# Dysregulation of cAMP-PKA-mediated mechanisms disrupts basal synaptic function in the absence of presynaptic FMRP

**DOI:** 10.64898/2026.10.01.755951

**Authors:** Simon Lecomte, Bruna Ferreira Mira, Noelle Grosjean, Christophe Mulle

## Abstract

The regulation of presynaptic mechanisms by the fragile X messenger ribonucleoprotein (FMRP) remains incompletely understood despite their potential importance to synaptic dysfunction reported in Fragile X syndrome. We combined post-developmental, cell-specific deletion of FMRP in dentate granule cells of the hippocampus with optogenetics, whole-cell patch-clamp electrophysiology and super resolution imaging. The selective loss of FMRP in DG cells did not alter their intrinsic excitability. The amplitude of synaptic currents was reduced at the output mossy fiber synapses in the absence of presynaptic FMRP, whereas short-term facilitation remained intact. We demonstrate that basal synaptic activity is regulated by PKA at mature mossy fiber synapses, and our results indicate alterations in cAMP-PKA signaling and a potential reduction in the readily releasable pool size in mossy fiber boutons lacking FMRP. Presynaptic FMRP is therefore required for efficient basal neurotransmission at mature mossy fiber synapses and likely acts through cAMP-PKA-dependent signaling pathway.

**Significance statement:** The fragile X messenger ribonucleoprotein (FMRP) has been extensively studied as a regulator of postsynaptic mechanisms. However, its physiological functions in presynaptic terminals remain incompletely understood. Using post-developmental, cell-specific deletion of FMRP, we show that presynaptic FMRP is required to maintain efficient basal neurotransmission at hippocampal mossy fiber synapses and identify cAMP-PKA signaling as a potential mechanism for this effect. Taken together, these results highlight a physiological role for presynaptic FMRP during basal synaptic activity and demonstrate how time-controlled gene-editing approaches via viral infection can distinguish primary molecular functions from developmental compensations, thereby shedding new light on both the presynaptic function of FMRP and the pathophysiology of Fragile X syndrome.

**Highlights:**

- Presynaptic loss of FMRP reduces basal synaptic activity at hippocampal Mf synapses.
- Basal synaptic activity is regulated by PKA in adult Mf synapses.
- AC-cAMP-PKA pathway is impaired during basal synaptic activity in the absence of presynaptic FMRP.
- RRP size is very likely reduced in AZs at Mf terminals.

## Introduction

Synaptic transmission is the fundamental mechanism by which neurons communicate and neuronal ensembles process information in the brain. Its precise regulation depends on the coordinated action of numerous signaling pathways that control neurotransmitter release, synaptic strength and activity-dependent plasticity (de Jong & Verhage, 2009; Kennedy, 2016; S. Li & Sheng, 2022; Saneyoshi et al., 2010). Alterations in these mechanisms are increasingly recognized as central contributors to neurodevelopmental disorders, including autism spectrum disorder and intellectual disability (Liao et al., 2026; Verpelli & Sala, 2012; Yan et al., 2016; Zhao et al., 2026; Zoghbi & Bear, 2012). The fragile X messenger ribonucleoprotein (FMRP) occupies a central position in proteins whose dysfunction contributes to neurodevelopmental disorders. Silencing of the *Fmr1* gene (located on the X chromosome) causes Fragile X syndrome (FXS), the most common inherited form of intellectual disability and a leading monogenic cause of autism spectrum disorder (Hagerman et al., 2017; Juárez et al., 2024). FMRP is an RNA- and ribosome-binding protein (Ashley et al., 1993; E. Chen et al., 2014) that controls the localization, stability and translation of hundreds of neuronal mRNAs in all subcellular compartments (Anderson et al., 2016; Ascano et al., 2012; Darnell et al., 2011; Klemmer et al., 2011; Maurin et al., 2018; Suhl et al., 2014). Understanding how FMRP shapes synaptic physiology is essential both for elucidating normal circuit function and for identifying the cellular mechanisms underlying the associated pathological conditions.

Over the past two decades, FMRP has emerged as a major regulator of neuronal excitability, connectivity and synapse development (Contractor et al., 2015; Hanson & Madison, 2007; Liu et al., 2022; Scharkowski et al., 2018). Most mechanistic studies have focused on the postsynaptic functions of FMRP. FMRP regulates dendritic protein synthesis, actin cytoskeleton remodeling, glutamate receptor expression and activity, as well as several intracellular signaling cascades, including PI3K-AKT-mTOR, ERK-MAPK, Wnt and calcium-dependent pathways (Brager & Johnston, 2014; Chmielewska et al., 2019; Corti & Duarte, 2025; Dictenberg et al., 2008; Gross et al., 2010; J. Huang et al., 2015; Huber et al., 2002; Luo et al., 2023; Meredith et al., 2007; Michaelsen-Preusse et al., 2018; Salcedo-Arellano et al., 2021; Thomazeau et al., 2021; Yang et al., 2019). These studies have established FMRP as a key postsynaptic regulator of synaptic function and plasticity.

Accumulating evidence indicates that FMRP also exerts important presynaptic functions (X. Wang et al., 2023). FMRP regulates axonal development and presynaptic local protein synthesis (Akins et al., 2009; Ivanco & Greenough, 2002; Monday et al., 2022; Till et al., 2011; Zimmer et al., 2017). Electrophysiological studies have further shown that FMRP influences action potential properties, presynaptic excitability, synaptic plasticity and activity-dependent bulk endocytosis (Bonnycastle et al., 2022; Deng et al., 2011, 2013, 2019; Deng & Klyachko, 2021; Klemmer et al., 2011; Monday et al., 2022; Myrick et al., 2015; Routh et al., 2017). However, despite growing recognition of these presynaptic functions, the role of presynaptic FMRP during basal neurotransmission remains unsettled (Patel et al., 2013; X.-S. Wang et al., 2014).

One signaling pathway repeatedly implicated in FXS is the cyclic AMP (cAMP) pathway. Since the pioneering work of Berry-Kravis and colleagues, numerous studies have reported reduced intracellular cAMP signaling following the loss of FMRP, leading to the proposal of the “cAMP theory” of FXS (Berry-Kravis et al., 1995; Berry-Kravis & Ciurlionis, 1998; Berry-Kravis & Huttenlocher, 1992; Berry-Kravis & Sklena, 1993; Kelley et al., 2007, 2008). More recent work has identified molecular links between FMRP and cAMP homeostasis. FMRP directly regulates transcripts encoding adenylate cyclase (AC) type 1 (Darnell et al., 2011; Maurin et al., 2018) and several phosphodiesterase subtypes. Therefore, the loss of FMRP increases phosphodiesterases expression (i.e. PDE2A, PDE4D and PDE10A) and disrupts cAMP-dependent signaling (Bardoni et al., 2025; Berry-Kravis et al., 2021; Choi et al., 2015; Darnell et al., 2011; Delhaye et al., 2024; Delhaye & Bardoni, 2021; Jonak et al., 2021; Maurin et al., 2018, 2019; Rosenheck et al., 2021; Schiavi et al., 2022). Although these observations strongly suggest an interaction between FMRP and cAMP signaling through PDEs, whether this interaction contributes directly in shaping presynaptic functions remains unknown.

Hippocampal mossy fiber (Mf) synapses provide a powerful system in which to address these questions. These giant glutamatergic boutons possess highly specialized presynaptic machinery and express remarkable forms of presynaptic facilitation (Marneffe et al., 2025). Synaptic plasticity, more particularly post-tetanic potentiation (PTP) and long-term potentiation (LTP), critically depend on presynaptic cAMP-PKA signaling (Bruentgens et al., 2026; Fukaya, Miyano, et al., 2023; Kim et al., 2024; Shahoha et al., 2022; Vandael et al., 2020; Weisskopf et al., 1994). Their dependence on presynaptic signaling pathways therefore makes Mf synapses an ideal model for investigating the physiological functions of presynaptic FMRP.

An additional challenge in understanding FMRP physiology is that most previous studies have relied on constitutive knockout and mosaic models. Although these models have provided fundamental insights into FXS pathophysiology, they inevitably combine the direct consequences of FMRP loss on neuronal function with the role of FMRP on the formation and maturation of neural circuits. Consequently, distinguishing the primary cell-autonomous physiological functions of FMRP from a developmental role and potential secondary compensatory mechanisms developed in its absence in the whole circuit remains difficult (Pfeiffer & Huber, 2009).

To address these questions, we selectively deleted FMRP from mature DG cells using a post-developmental, cell-specific viral strategy combined with optogenetics, whole-cell patch-clamp electrophysiology at hippocampal Mf-CA3 synapses and super resolution imaging of Mf boutons. These approaches allowed us to investigate the presynaptic functions of FMRP while minimizing developmental confounds. Our findings reveal an important role for presynaptic FMRP in maintaining efficient basal neurotransmission and identify the cAMP-PKA signaling pathway as a candidate mechanism underlying this regulation.

## Results

### Deletion of FMRP does not affect DG cells excitability

FMRP is known to interact either physically or indirectly (through its translational role) with ion channels in neurons (Deng & Klyachko, 2021) and thus to impact neuronal excitability (Contractor et al., 2015; Liu et al., 2022). Therefore, some studies reported an alteration of subthreshold membrane properties of neurons lacking FMRP, such as resting membrane potential (M. Shen et al., 2023) or input resistance (Luque et al., 2017; M. Shen et al., 2023), and also of suprathreshold membrane properties, such as rheobase (Deng et al., 2024), spiking rate (Deng et al., 2024; Routh et al., 2017; Zhang et al., 2014), firing threshold (Deng et al., 2024; Deng & Klyachko, 2016), action potential (AP) amplitude (Routh et al., 2017), duration (Deng et al., 2013; Luque et al., 2017; Myrick et al., 2015; Routh et al., 2017), afterhyperpolarization (AHP) and afterdepolarization (ADP) amplitudes (Luque et al., 2017). However, depending on the species, strains, age, brain region, cell subtype, postsynaptic cell-target or even subcellular compartment, the cellular mechanisms altered by loss of FMRP differ markedly between studies, and at times appear contradictory. Hence it is important to assess whether deletion of FMRP in DG cells after their maturation has an impact on their ability to fire action potentials and on the action potential properties themselves.

We first confirmed that our viral approach enabled the deletion of FMRP in DG cells of *Fmr1^flox/y^* mice after stereotaxic injection of the lentivirus LV.C1qL2.ChIEF-Tom-2A-Cre (Barthet et al., 2018) (**Figure 1A-B**). Indeed, Cre^+^ cells in the DG no longer expressed the protein FMRP, in contrast to Cre^−^cells (**Figure 1D**). We then recorded from DG cells expressing the viral construct in WT vs. *Fmr1^flox/y^* mice and checked for their membrane properties (**Figure 1C**). We also recorded DG cells not expressing the virus (non-infected, NI cells) regardless of their position within the granule cell layer, to have a control of the effect of the virus expression on neuronal excitability. We used 500 ms steps of hyperpolarizing and depolarizing currents (**Figure 1E**). The RMP of neurons (NI: -69.3 ± 2.3, *n*=13; WT: -66 ± 5.7, *n*=20; cKO: -68.1 ± 2.1, *n*=15; NI vs. WT: *p*=0.0718; NI vs. cKO: *p*=0.3638; WT vs. cKO: *p*=0.3571; *W*(2, 29.4)=3, *p*=0.063; **Figure 1F**), input resistance (NI: 425.4 ± 120.4, *n*=13; WT: 401 ± 116.6, *n*=20; cKO: 395.5 ± 70.9, *n*=15; NI vs. WT: *p*=0.9165; NI vs. cKO: *p*=0.817; WT vs. cKO: *p*=0.9974; *W*(2, 26.8)=0.3, *p*=0.7418; **Figure 1G**), rheobase (NI: 75.4 ± 44.8, *n*=13; WT: 65 ± 23.3, *n*=20; cKO: 64 ± 18.8, *n*=15; NI vs. WT, NI vs. cKO, WT vs. cKO, *p*>0.9999; *H*(2)=0.08, *p*=0.9605; **Figure 1H**), and spiking rate (NI: *n*=13, WT: *n*=20, cKO: *n*=15; *F*(2, 46)=1, *p*=0.3731; **Figure 1I**) were not statistically different between WT and FMRP-deficient cells. Therefore, it can be concluded that DG cells lacking FMRP showed the same profile of excitability as FMRP-containing DG cells. We then analyzed the AP properties (**Figure 1J**). The firing threshold (NI: -32.4 ± 5.1, *n*=13; WT: -35.6 ± 6.3, *n*=20; cKO: -35.9 ± 9.4, *n*=15; NI vs. WT: *p*=0.6032; NI vs. cKO: *p*=0.2346; WT vs. cKO: *p*>0.9999; *H*(2)=3.2, *p*=0.2005; **Figure 1K**), full width at half maximum (FWHM) (NI: 1.9 ± 0.2, *n*=13; WT: 1.9 ± 0.4, *n*=20; cKO: 1.8 ± 0.3, *n*=15; NI vs. WT: *p*=0.9986; NI vs. cKO: *p*=0.9236; WT vs. cKO: *p*=0.9833; *W*(2, 29.9)=0.2, *p*=0.8575; **Figure 1M**) and AHP (NI: 19.6 ± 3.9, *n*=13; WT: 15.2 ± 3.8, *n*=20; cKO: 15.7 ± 3.4, *n*=15; NI vs. WT: *p*=0.0203; NI vs. cKO: *p*=0.0308; WT vs. cKO: *p*>0.9999; *H*(2)=8.9, *p*=0.0117; **Figure 1N**) were similar between action potentials from WT and FMRP-deficient DG cells. However, the AP amplitude (NI: 81.2 ± 9.1, *n*=13; WT: 84.9 ± 6.9, *n*=20; cKO: 91.9 ± 8.1, *n*=15; NI vs. WT: *p*>0.9999; NI vs. cKO: *p*=0.0026; WT vs. cKO: *p*=0.0125; *H*(2)=13, *p*=0.0016; **Figure 1L**), was significantly increased in FMRP-deficient DG cells. This effect is consistent with the literature and it is likely caused by an increase in voltage-gated sodium conductance due to indirect interaction between FMRP and voltage-gated sodium channels at the soma and axonal initial segment (Deng & Klyachko, 2016, 2021; Routh et al., 2017).

**Figure 1.**
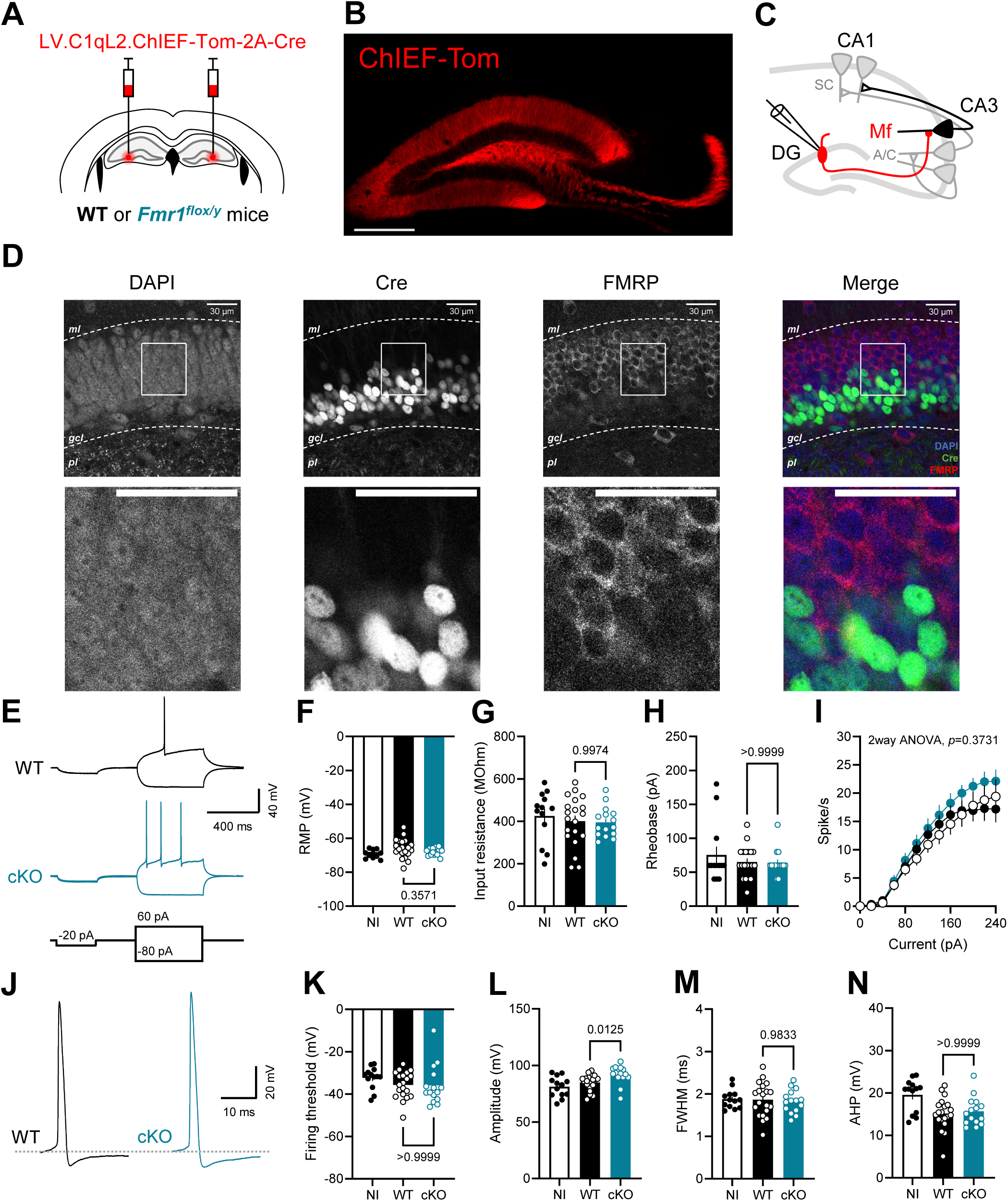
Post-developmental and selective deletion of FMRP in dentate granule cells does not alter their intrinsic excitability. (A) Illustration of stereotaxic injection leading to expression of ChIEF and Cre in the dentate gyrus of P25-P30 wild-type (WT, black) and dentate gyrus-specific *Fmr1* conditional knock-out (cKO, blue) mice. (B) Image of ChIEF expression selectively in the dentate gyrus 4 weeks after stereotaxic injection. Scale bar: 300 µm. (C) Scheme of acute hippocampal slice with a recording pipette in the soma of a dentate granule cell to record membrane properties. A/C: associative/commissural loop; CA1-3: cornu ammonis subfields; DG: dentate gyrus; Mf: mossy fiber pathway; SC: Schaffer collateral pathway. (D) *Top panels*: Representative immunofluorescence images showing DAPI, Cre, and FMRP expression in the soma of dentate granule cells from one FMRP DG-cKO mouse. *Bottom panels*: Close-up of boxed areas showing the absence of FMRP expression in Cre^+^ granule cells and preserved FMRP signal in non-recombined (Cre^−^) granule cells. *gcl*: granule cell layer; *ml*: molecular layer; *pl*: polymorphic layer. Scale bar: 30 μm. (E) Representative recordings of 500 milliseconds steps of current (hyperpolarizing and depolarizing) from dentate granule cells in acute brain slices with a 300 ms hyperpolarizing pre-step of -20 pA. (F) The resting membrane potential is not affected by the lack of FMRP (Welch’s ANOVA, *W*(2, 29.4)=3; Dunnett’s post-hoc test, *p*=0.3571). (G) The input resistance is not affected by the lack of FMRP (Welch’s ANOVA, *W*(2, 26.8)=0.3; Dunnett’s post-hoc test, *p*=0.9974). (H) The rheobase is not changed by the lack of FMRP (Kruskal-Wallis test, *H*(2)=0.08; Dunn’s T3 post-hoc test, *p*>0.9999). (I) The spike rate is not altered by the lack of FMRP (2way ANOVA, main effect between groups, *F*(2, 46)=1, *p*=0.3731). (J) Representative traces of action potentials obtained from the rheobase current step of dentate granule cells in acute brain slices. (K) The firing threshold is not affected by the lack of FMRP (Kruskal-Wallis test, *H*(2)=3.2; Dunn’s T3 post-hoc test, *p*>0.9999). (L) The amplitude of action potential is increased in the absence of FMRP (Kruskal-Wallis test, *H*(2)=13; Dunn’s T3 post-hoc test, *p*=0.0125). (M) The full-width at half maximum is not changed by the lack of FMRP (Welch’s ANOVA, *W*(2, 29.9)=0.2; Dunnett’s post-hoc test, *p*=0.9833). (N) The afterhyperpolarization phase of the action potential is not altered by the lack of FMRP (Kruskal-Wallis test, *H*(2)=8.9; Dunn’s T3 post-hoc test, *p*>0.9999). NI: *n*=13 cells; WT: *n*=20 cells; cKO: *n*=15 cells. NI: *N*=8 mice; WT: *N*=7 mice; cKO: *N*=5 mice. Data are shown as mean ± SEM.

The viral construct used allowed us to selectively stimulate FMRP-deficient Mfs when expressed in *Fmr1^flox/y^* mice under blue light illumination (1 ms pulses at 470 nm) to study the impact of presynaptic FMRP deletion on synaptic activity. It was important to determine whether the properties of APs observed during current steps was maintained during optogenetic stimulations (**Figure S1A**), in order to ensure the reliability of this stimulation and to allow for a valid comparison of postsynaptic currents between experimental groups. Input-output curve of AP probability (%) as a function of LED irradiance at the focal plane showed that at 11.36 mW/mm², light pulses always caused an action potential in both groups (**Figure S1B**). For this reason, an irradiance of 11.36 mW/mm² was chosen as the stimulation parameter during all the study and consequently, for the following analysis of AP properties. In DG cells, the firing threshold (WT: -40.3 ± 6.4, *n*=21; cKO: -41.9 ± 4.4, *n*=20; *t*(35.4)=0.9, *p*=0.3565; **Figure S1C**), FWHM (WT: 1.5 ± 0.2, *n*=21; cKO: 1.7 ± 0.3, *n*=20; *t*(37.6)=1.6, *p*=0.1129; **Figure S1E**), AHP (WT: 10.4 ± 5.8, *n*=21; cKO: 10.9 ± 5.4, *n*=20; *U*=191, *p*=0.633; **Figure S1F**) and ADP (WT: 10.4 ± 1.9, *n*=21; cKO: 9.2 ± 3.3, *n*=20; *t*(30.4)=1.5, *p*=0.1541; **Figure S1G**) of light-evoked APs were not statistically different between WT and FMRP-deficient DG cells. Surprisingly, the amplitude of APs elicited by optogenetic stimulation was similar between groups (WT: 88.7 ± 5.1, *n*=21; cKO: 88.4 ± 9.3, *n*=20; *U*=187, *p*=0.5612; **Figure S1D**). From these data it can be concluded that the optogenetic stimulation was reliable and rigorously similar in both groups (for all the parameters analyzed). Therefore, any differences in postsynaptic currents between groups could not be imputed to stimulation artifacts (i.e. taller or broader action potentials).

### Impaired synaptic amplitude in FMRP DG-cKO synapses depends on presynaptic mechanisms

The role of presynaptic FMRP during basal synaptic transmission remains contradictory. One study conducted on mosaic mice (Patel et al., 2013), demonstrated that deletion of presynaptic FMRP impaired excitatory transmission onto fast-spiking interneurons in layers 4-5 of the mouse barrel cortex under basal conditions. In contrast, no synaptic deficit was observed at excitatory-to-excitatory synapses, suggesting that the contribution of presynaptic FMRP may depend on the postsynaptic cell target.

To test whether presynaptic FMRP regulates Mf-CA3 synaptic properties, we patched CA3 neurons from control (injected WT mice) and FMRP DG-cKO mice (injected *Fmr1^flox/y^*mice) and recorded excitatory postsynaptic currents in CA3 pyramidal neurons evoked by 1 ms light pulses at 0.1Hz focused on the DG (**Figure 2A-B**). Mf-EPSCs amplitude (WT: 111.3 ± 79.9, *n*=68; cKO: 54.4 ± 41.2, *n*=75; *U*=1076, *p*<0.0001; **Figure 2C**, **H**, **Table 1**) and Mf-EPSCs potency (WT: 123.7 ± 75.8, *n*=68; cKO: 73.1 ± 38.2, *n*=75; *U*=1156, *p*<0.0001; **Figure 2D**, **G**, **Table 2**) were decreased in the absence of presynaptic FMRP. Failure rate (WT: 14.1 ± 14, *n*=68; cKO: 30.5 ± 18.6, *n*=75; *U*=1207, *p*<0.0001; **Figure 2E**) and CV (WT: 75.1 ± 26.8, *n*=68; cKO: 102.6 ± 36, *n*=75; *U*=1384, *p*<0.0001; **Figure 2F**) were increased in the absence of presynaptic FMRP. Together, these findings indicate that presynaptic FMRP contributes to maintaining robust and reliable basal excitatory transmission on excitatory neurons at the Mf-CA3 synapse.

**Figure 2.**
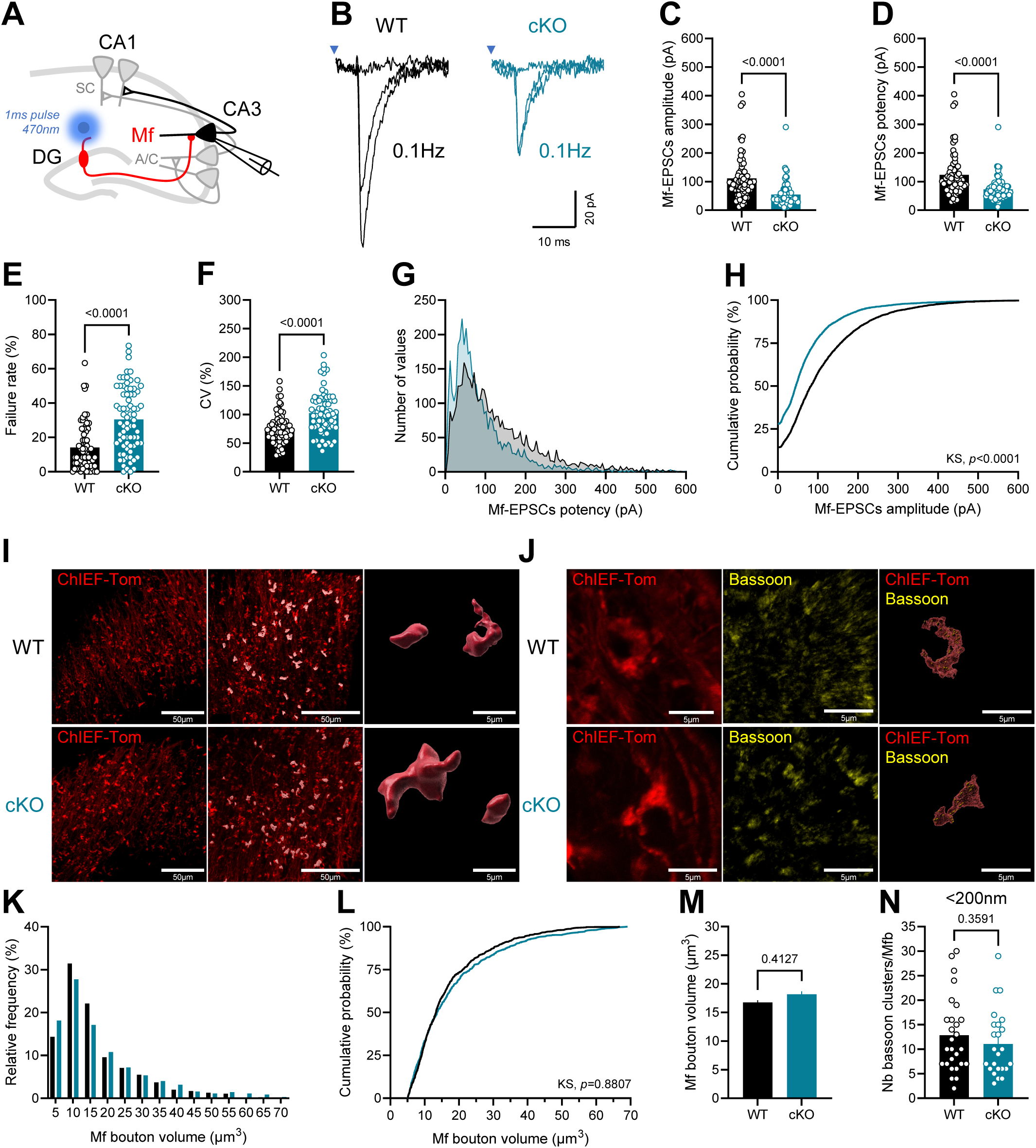
Basal Mf transmission is reduced in the absence of presynaptic FMRP while the volume of Mf boutons and their number of active zones containing bassoon remain unchanged. (A) Scheme of acute hippocampal slice with a blue light pulse (1 ms, 470 nm) in the dendrites of DG cells to elicit action potentials and a recording pipette in the soma of a CA3 pyramidal cell to record Mf synaptic currents. A/C: associative/commissural loop; CA1-3: cornu ammonis subfields; DG: dentate gyrus; Mf: mossy fiber pathway; SC: Schaffer collateral pathway. (B) Representative traces of Mf-EPSCs obtained during a 0.1 Hz stimulation protocol from control (black) and FMRP DG-cKO (blue) mice. (C) Mean amplitude of Mf-EPSCs is decreased in the absence of presynaptic FMRP (Mann-Whitney test, *U*=1076, *p*<0.0001). (D) Mean potency of MF-EPSCs (failures are removed from the analysis) is decreased in the absence of presynaptic FMRP (Mann-Whitney test, *U*=1156, *p*<0.0001). (E) Mean failure rate of Mf-EPSCs is increased in the absence of presynaptic FMRP (Mann-Whitney test, *U*=1207, *p*<0.0001). (F) Mean coefficient of variation of Mf-EPSCs is increased in the absence of presynaptic FMRP (Mann-Whitney test, *U*=1384, *p*<0.0001). WT: *n*=68 cells; cKO: *n*=75 cells. WT: *N*=49 mice; cKO: *N*=58 mice. (G) Distribution of Mf-EPSCs potency (0 to 600 pA) from control and FMRP DG-cKO mice. Bins of 5 pA. WT: *n*=4350 Mf-EPSCs; cKO: *n*=3975 Mf-EPSCs. WT: *N*=68 cells; cKO: *N*=75 cells. (H) Cumulative probability analysis of the distribution of Mf-EPSCs amplitude shows a decrease of Mf-EPSCs amplitude in the absence of presynaptic FMRP (Kolmogorov-Smirnov test, *D*=0.2, *p*<0.0001). No bins. WT: *n*=5184 Mf-EPSCs; cKO: *n*=5582 Mf-EPSCs. WT: *N*=68 cells; cKO: *N*=75 cells. (I) *Left panels*: Representative images showing ChIEF and tdTomato expression in the Mf boutons from control and FMRP DG-cKO mice. Scale bar: 50 µm. *Middle panels*: Imaris detection and isolation of Mf boutons. Scale bar: 50 µm. *Right panels*: Imaris 3D reconstruction of Mf boutons. Scale bar: 5 µm. (J) *Left panels*: Representative immunofluorescence images showing ChIEF and tdTomato expression in the Mf boutons from control and FMRP DG-cKO mice. Scale bar: 5 µm. *Middle panels*: Representative immunofluorescence images showing bassoon expression in Mf boutons. Scale bar: 5 µm. *Right panels*: Imaris 3D reconstruction of Mf boutons surface together with 3D reconstructed bassoon clusters. Scale bar: 5 µm. (K) Histograms of the distribution of Mf boutons volume. Bins of 5 µm^3^. (L) Cumulative probability analysis of the distribution of Mf boutons volume shows no statistical difference in the volume of Mf boutons between control and FMRP DG-cKO mice (Kolmogorov-Smirnov test, *D*=0.03, *p*=0.8807). No bins. (M) Mean volume of Mf boutons is not altered by the lack of FMRP (Mann-Whitney test, *U*=283795, *p*=0.4127). WT: *n*=837 Mf boutons; cKO: *n*=695 Mf boutons. WT: *N*=3 mice; cKO: *N*=3 mice. (N) Mean number of bassoon clusters per Mf bouton within 200 nm from plasma membrane is not changed by the absence of FMRP (Mann-Whitney test, *U*=275, *p*=0.3591). WT: *n*=27 Mf boutons; cKO: *n*=24 Mf boutons. WT: *N*=3 mice; cKO: *N*=3 mice.

**Table 1.** Distribution of Mf-EPSCs amplitude recorded at 0.1 Hz (including failures).

| Parameters | WT | cKO |
| --- | --- | --- |
| Mode | 0 pA | 0 pA |
| Mean | 110.2 pA | 66.6 pA |
| Median | 82.5 pA | 44.9 pA |
| SD | 106.1 pA | 86.4 pA |
| CV | 96.3 % | 128.2 % |
| Number of EPSCs | 5184 | 5582 |
| Number of cells | 68 | 75 |

**Table 2.** Distribution of Mf-EPSCs potency recorded at 0.1 Hz (without failures).

| Parameters | WT | cKO |
| --- | --- | --- |
| Mode | 47 pA | 40 pA |
| Mean | 130.3 pA | 92.4 pA |
| Median | 100.3 pA | 66.2 pA |
| SD | 103.7 pA | 87.4 pA |
| CV | 79.6 % | 94.6 % |
| Number of EPSCs | 4350 | 3975 |
| Number of cells | 68 | 75 |

Presynaptic alterations can have a significant impact on synaptic transmission through postsynaptic mechanisms, known as “transsynaptic mechanisms” (Chia et al., 2013; K. Shen & Scheiffele, 2010; Verpoort & de Wit, 2024). When reporting changes in synaptic activity, it is important to examine the synaptic locus responsible for these changes. Mean postsynaptic current amplitude is defined by three quantal parameters (Johnson & Wernig, 1971; Korn & Faber, 1991; McLachlan, 1978; Redman, 1990). Namely, the mean number of independent and active release sites ⟨n⟩, their mean release probability following an incoming AP ⟨p⟩ and the mean postsynaptic response elicited by one quantal release event ⟨q⟩. From the standard Katz model of synaptic transmission these parameters are associated with a synaptic locus: ⟨n⟩ and ⟨p⟩ are presynaptic parameters and ⟨q⟩ is mainly postsynaptic.

To test whether postsynaptic mechanisms could explain the alteration of basal synaptic transmission in the absence of presynaptic FMRP, we bath applied cadmium (Cd^2+^) and recorded the amplitude of quantal synaptic events (see **Figure 6A-B**). Both groups showed similar amplitudes of quantal events (WT: 37.3 ± 16.1, *n*=8; cKO: 41.2 ± 22.4, *n*=8; *U*=29, *p*=0.7984; **Figure 6C**). This result suggests that presynaptic loss of FMRP affects presynaptic function at Mf synapses. A convergent observation is that amplitude of Mf-EPSCs was reduced in the absence of presynaptic FMRP together with an increase in coefficient of variation (**Figure 2C, F**), covariation of these two parameters is linked to presynaptic mechanisms change. In order to specify which presynaptic parameter is involved (i.e. ⟨n⟩ or ⟨p⟩) we first looked at an often-used proxy of initial ⟨p⟩: the paired-pulse ratio (PPR). PPR was similar in both groups (WT: 2.7 ± 1.4, *n*=25; cKO: 2.7 ± 2.2, *n*=26; *U*=241, *p*=0.1162; **Figure 3I**). This suggests that rather than the probability of release ⟨p⟩, the number of independent and active release sites is affected in the absence of presynaptic FMRP. In line with this hypothesis, the failure rate was increased in the absence of presynaptic FMRP (**Figure 2E**), which can be an indication of a defect in the number of active release sites. Mfs possess giant presynaptic boutons containing up to ∼45 active zones (AZs) (Chicurel & Harris, 1992; Rollenhagen et al., 2007) with ∼40 vesicles inside the readily releasable pool (RRP) per AZ (Maus et al., 2020; Vandael & Jonas, 2024). Due to their large number of active zones, a change in the number of independent and active release sites ⟨n⟩ can be due either to a change in the number of active zones per bouton, a change in number of docked/primed vesicles inside the RRP or both. To decipher which mechanism is involved we used confocal and STED microscopy, and measured both the volume of Mf boutons (**Figure 2I**) as well as the number of active zones (i.e. bassoon clusters) per bouton (**Figure 2J**). Both the mean volume of Mf boutons (WT: 16.8 ± 11, *n*=837; cKO: 18.2 ± 13.3, *n*=695; *U*=283795, *p*=0.4127; **Figure 2K-M**) and the mean number of bassoon clusters per bouton (WT: 12.9 ± 7.7, *n*=27; cKO: 11.1 ± 6.7, *n*=24; *U*=275, *p*=0.3591; **Figure 2N**) were similar in both groups. These results suggest that presynaptic FMRP does not affect the number of AZs containing bassoon at Mf boutons; rather, it likely plays a role in determining the number of active release sites, hence the size of their RRP.

**Figure 3.**
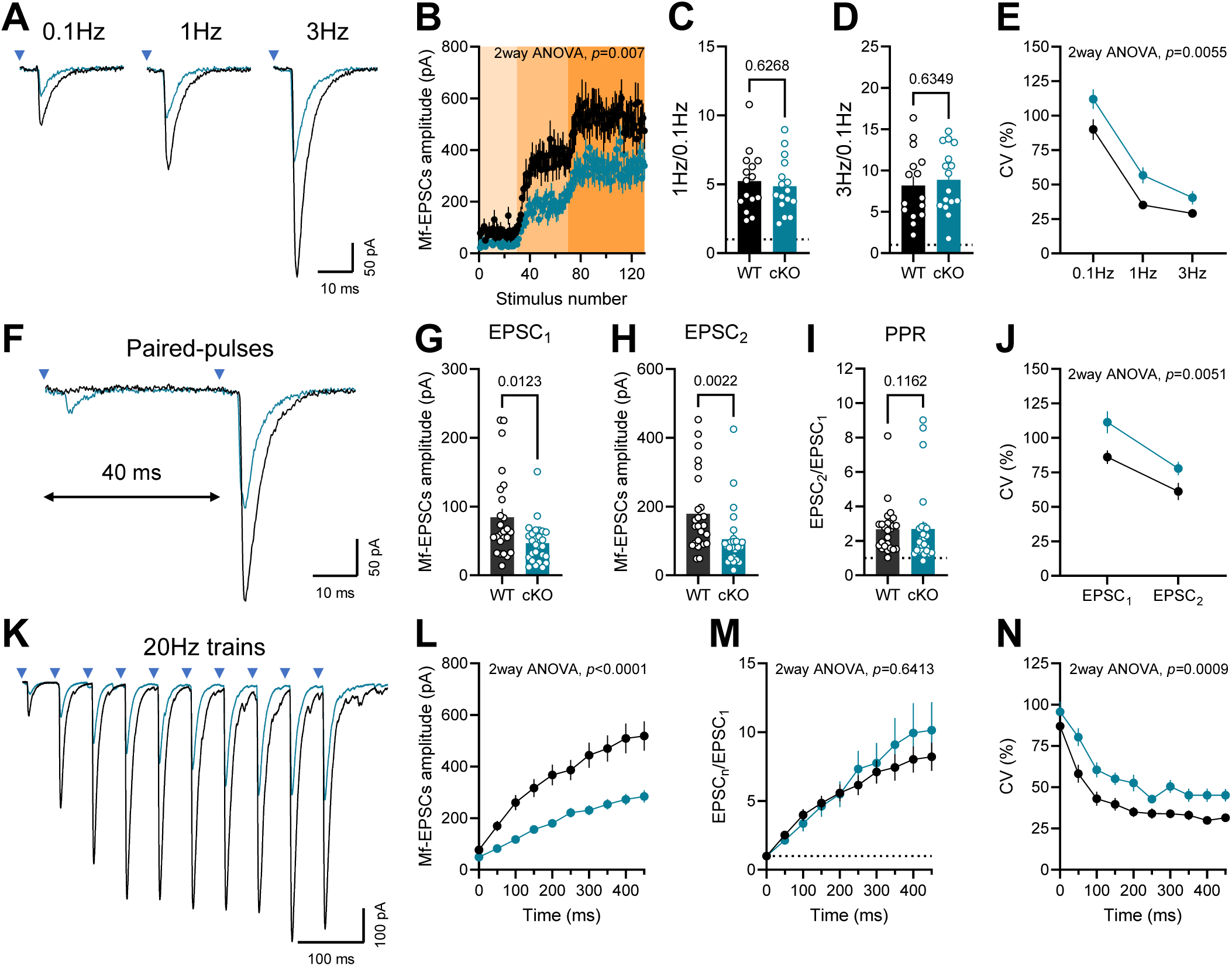
Magnitude of mossy fiber facilitation is unaltered in the absence of presynaptic FMRP. (A) Representative traces of Mf-EPSCs obtained during 0.1 Hz, 1 Hz and 3 Hz protocols from CA3 pyramidal neurons in acute brain slices from control (black) and FMRP DG-cKO (blue) mice. (B) Time course of Mf-EPSCs amplitude at 0.1 Hz (light orange), 1 Hz (orange) and 3 Hz (dark orange). Amplitudes of Mf-EPSCs are reduced in the absence of presynaptic FMRP (2way ANOVA, main effect between groups, *F*(1, 29)=8.4, *p*=0.007). (C) Frequency facilitation at 1 Hz (ratio between mean amplitude at 1 Hz and at 0.1 Hz) is not affected by the lack of presynaptic FMRP (Welch’s t-test, *t*(28)=0.5, *p*=0.6268). (D) Mean frequency facilitation at 3 Hz (ratio between mean amplitude at 3 Hz and at 0.1 Hz) is not affected by the lack of presynaptic FMRP (Welch’s t-test, *t*(28.4)=0.5, *p*=0.6349). (E) Mean coefficient of variation of Mf-EPSCs during 0.1 Hz, 1 Hz and 3 Hz is increased in the absence of presynaptic FMRP (2way ANOVA, main effect between groups, *F*(1, 29)=9, *p*=0.0055). WT: *n*=15 cells; cKO: *n*=16 cells. WT: *N*=8 mice; cKO: *N*=10 mice. (F) Representative traces of Mf-EPSCs obtained during a paired-pulses protocol with an inter-event interval of 40 ms from CA3 pyramidal neurons in acute brain slices from control and FMRP DG-cKO mice. (G) Mean amplitude of the first Mf-EPSCs is decreased in the absence of presynaptic FMRP during a paired-pulses protocol (Mann-Whitney test, *U*=193, *p*=0.0123). (H) Mean amplitude of the second Mf-EPSCs is decreased in the absence of presynaptic FMRP during a paired-pulses protocol (Mann-Whitney test, *U*=165, *p*=0.0022). (I) Paired-pulses ratio (ratio between mean amplitude of the second pulse and the first) is not affected by the lack of presynaptic FMRP (Mann-Whitney test, *U*=241, *p*=0.1162). (J) Mean coefficient of variation of Mf-EPSCs during a paired-pulses protocol is increased in the absence of presynaptic FMRP (2way ANOVA, main effect between groups, *F*(1, 49)=8.6, *p*=0.0051). WT: *n*=25 cells; cKO: *n*=26 cells. WT: *N*=18 mice; cKO: *N*=18 mice. (K) Representative traces of Mf-EPSCs obtained during a train of 10 light pulses at 20 Hz from CA3 pyramidal neurons in acute brain slices from control and FMRP DG-cKO mice. (L) Mean amplitudes of Mf-EPSCs within the train are decreased in the absence of presynaptic FMRP (2way ANOVA, main effect between groups, *F*(1, 62)=19.2, *p*<0.0001). (M) Train facilitation (ratio between each Mf-EPSCs within the train and the first one) is not affected by the lack of presynaptic FMRP (2way ANOVA, main effect between groups, *F*(1, 62)=0.2, *p*=0.6413). (N) Mean coefficient of variation of Mf-EPSCs across the train is increased in the absence of presynaptic FMRP (2way ANOVA, main effect between groups, *F*(1, 62)=12.1, *p*=0.0009). WT: *n*=31 cells; cKO: *n*=33 cells. WT: *N*=20 mice; cKO: *N*=21 mice. Data are shown as mean ± SEM.

### Presynaptic loss of FMRP does not affect short-term facilitation at Mf synapses

We examined the role of presynaptic FMRP in the pronounced forms of short-term plasticity at Mf-CA3 synapses (Marneffe et al., 2025). We first checked frequency facilitation which consists in increasing frequency stimulation from 0.1 Hz to 1 Hz and 3 Hz; **Figure 3A**), paired-pulse facilitation which consists in paired stimulations with 40 ms interval (**Figure 3F**) and train facilitation which consists in a 10 stimulations train at 20 Hz (**Figure 3K**). In these experiments we repeatedly observed a consistent decrease in the amplitude of Mf-EPSCs in the absence of presynaptic FMRP, during frequency facilitation (WT: *n*=15, cKO: *n*=16; *F*(1, 29)=8.4, *p*=0.007; **Figure 3B**), during paired-pulse facilitation for both the first EPSC (WT: 84.5 ± 61.1, *n*=25; cKO: 46.7 ± 29.6, *n*=26; *U*=193, *p*=0.0123; **Figure 3G**) and the second EPSC (WT: 179 ± 115.5, *n*=25; cKO: 105 ± 84.3, *n*=26; *U*=165, *p*=0.0022; **Figure 3H**), and finally during train facilitation (WT: *n*=31, cKO: *n*=33; *F*(1, 62)=19.2, *p*<0.0001; **Figure 3L**).

The magnitude of facilitation was similar in both groups: for frequency facilitation at 1 Hz (WT: 5.2 ± 2.2, *n*=15; cKO: 4.9 ± 2, *n*=16; *t*(28)=0.5, *p*=0.6268; **Figure 3C**), for frequency facilitation at 3 HZ (WT: 8.2 ± 4.2, *n*=15; cKO: 8.9 ± 3.9, *n*=16; *t*(28.4)=0.5, *p*= 0.6349; **Figure 3D**), for paired-pulse ratios (**Figure 3I**) and for train facilitation (WT: *n*=31, cKO: *n*=33; *F*(1, 62)=0.2, *p*=0.6413; **Figure 3M**). The coefficient of variation was increased in the absence of presynaptic FMRP during frequency facilitation (WT: *n*=15, cKO: *n*=16; *F*(1, 29)=9, *p*=0.0055; **Figure 3E**), paired-pulse facilitation (WT: *n*=25, cKO: *n*=26; *F*(1, 49)=8.6, *p*=0.0051; **Figure 3J**), and train facilitation (WT: *n*=31, cKO: *n*=33; *F*(1, 62)=12.1, *p*=0.0009; **Figure 3N**).

Overall, we observed that: (1) the amplitude of Mf-EPSCs was reduced (**Figure 3B, G, H, L**), (2) the magnitude of Mf short-term facilitation was unaltered (**Figure 3C, D, I, M**) and (3) the coefficient of variation was increased in the absence of presynaptic FMRP (**Figure 3E, J, N**). These results show that the mechanisms of frequency facilitation and train facilitation are not altered by the loss of presynaptic FMRP at Mf synapses.

### Presynaptic loss of FMRP increases the magnitude of Mf-optoPTP

Another type of short-term facilitation is observed at Mf synapses: post-tetanic potentiation (PTP). While 3 trains of 100 electrical stimulations at 100 Hz induces a presynaptic form of LTP at Mfs (Monday et al., 2018; Monday & Castillo, 2017; Nicoll & Schmitz, 2005), only one train instead causes short-term plasticity (Griffith, 1990). FMRP has been shown to be crucial for LTP maintenance through its role in local protein synthesis at Mf terminals (Monday et al., 2022). However, its role during PTP has not been addressed. One major limitation of optogenetics is the frequency of stimulation channelrhodopsin variants can support. ChIEF (Chimera EF with I170V mutation) provides reliable stimulation only up to 25 Hz (Lin, 2011). Therefore, we used 20 Hz as the stimulation frequency to induce PTP with optogenetics (hence ‘optoPTP’) with a train of 50 pulses (**Figure 4A**).

**Figure 4.**
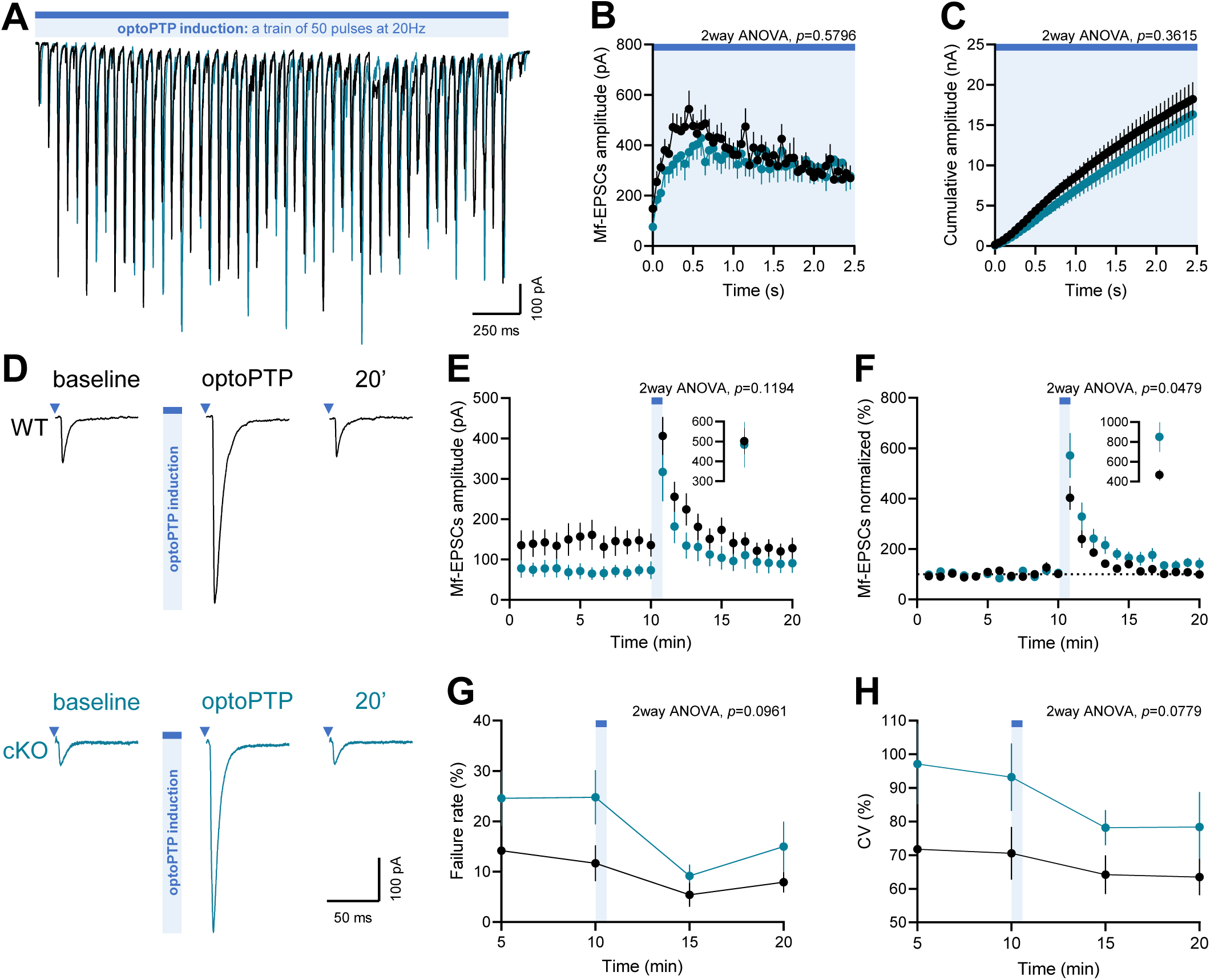
Optogenetically-induced PTP magnitude is increased with preserved absolute potentiation in the absence of presynaptic FMRP. (A) Representative traces of Mf-EPSCs recorded during optoPTP induction: a single train of 50 light pulses at 20 Hz (blue bar). Recordings from CA3 pyramidal neurons in acute brain slices from control (black) and FMRP DG-cKO (blue) mice. (B) Mean amplitudes of Mf-EPSCs within the train are not affected in the absence of presynaptic FMRP (2way ANOVA, main effect between groups, *F*(1, 49)=0.3, *p*=0.5796). (C) Cumulative amplitudes of Mf-EPSCs within the train are not affected in the absence of presynaptic FMRP (2way ANOVA, main effect between groups, *F*(1, 49)=0.8, *p*=0.3615). WT: *n*=23 cells; cKO: *n*=28 cells. WT: *N*=14 mice; cKO: *N*=20 mice. (D) Representative traces of Mf-EPSCs recorded at 0.1 Hz during 10 min baseline, 5 seconds after the optoPTP induction (optoPTP) and 10 min after optoPTP induction. Recordings from CA3 pyramidal neurons in acute brain slices from control and FMRP DG-cKO mice. (E) Time course of Mf-EPSCs amplitude for the experiment in (D) shows similar amplitudes in the absence of presynaptic FMRP (2way ANOVA, main effect between groups, *F*(1, 30)=2.6, *p*=0.1194). Time points are made by doing the average amplitude of 5 Mf-EPSCs. *Insert*: average amplitude of the first Mf-EPSC recorded 5 seconds after induction, shows similar absolute potentiation (Mann-Whitney test, *U*=106, *p*=0.423). (F) Time course of normalized Mf-EPSCs amplitude for the experiment in (D) shows increased amplitudes in the absence of presynaptic FMRP (2way ANOVA, main effect between groups, *F*(1, 30)=4.3, *p*=0.0479). Time points are made by doing the average amplitude of 5 Mf-EPSCs. *Insert*: average amplitude of the first Mf-EPSC recorded 5 seconds after induction, shows increased optoPTP magnitude in the absence of presynaptic FMRP (Mann-Whitney test, *U*=71, *p*=0.0318). (G) Time course of failure rate of Mf-EPSCs for the experiment in (D) shows similar failure rates in the absence of presynaptic FMRP (2way ANOVA, main effect between groups, *F*(1, 30)=3, *p*=0.0961). Time points are made by calculating failure rate during 5 min of recording. (H) Time course of coefficient of variation of Mf-EPSCs for the experiment in (D) shows similar coefficient of variation in the absence of presynaptic FMRP (2way ANOVA, main effect between groups, *F*(1, 30)=3.3, *p*=0.0779). Time points are made by calculating coefficient of variation during 5 min of recording. WT: *n*=16 cells; cKO: *n*=16 cells. WT: *N*=12 mice; cKO: *N*=13 mice. Data are shown as mean ± SEM.

Neither the amplitude of Mf-EPSCs within the induction train (WT: *n*=23, cKO: *n*=28; *F*(1, 49)=0.3, *p*=0.5796; **Figure 4B**) nor the cumulative amplitude across the train (WT: *n*=23, cKO: *n*=28; *F*(1, 49)=0.8, *p*=0.3615; **Figure 4C**) differed significantly between control and FMRP DG-cKO mice. This indicates that overall optoPTP induction is comparable in both groups. However, closer inspection of individual Mf-EPSCs reveals a reduction in the amplitude during the first 10 Mf-EPSCs in FMRP DG-cKO mice compared to control mice (consistent with the results described in **Figure 3L**) whereas no such difference was observed for the remaining part of the train. Mf-EPSCs were recorded at 0.1 Hz for 10 minutes before and 10 minutes after optoPTP induction (**Figure 4D**). First, the absolute amplitude of Mf-EPSCs during the baseline is significantly lower in FMRP DG-cKO mice compared to control mice (WT: 143.5 ± 128.1, *n*=16; cKO: 72.1 ± 71.9, *n*=16; *U*=74, *p*=0.0426; **Figure 4E**). However, the amplitude of the first Mf-EPSC recorded 5 seconds after induction does not differ between groups (WT: 501.2 ± 263.4, *n*=16; cKO: 483.9 ± 455.9, *n*=16; *U*=106, *p*=0.423; **Figure 4E**, insert). Hence, when amplitudes are normalized to baseline, FMRP DG-cKO mice display a significantly larger magnitude of potentiation just after optoPTP induction compared to control mice (WT: 466.6 ± 218, *n*=16; cKO: 850.2 ± 601.9, *n*=16; *U*=71, *p*=0.0318; **Figure 4F**, insert). Finally, the failure rate (WT: *n*=16, cKO: *n*=16; *F*(1, 30)=3, *p*=0.0961; **Figure 4G**) and coefficient of variation (WT: *n*=16, cKO: *n*=16; *F*(1, 30)=3.3, *p*=0.0779; **Figure 4H**) are not statistically different after optoPTP induction between FMRP DG-cKO and control mice.

Altogether these data support the idea that Mf-optoPTP is able to counteract the aforementioned phenotype induced by presynaptic FMRP deletion, namely reduced Mf-EPSCs amplitude during basal synaptic activity, increased failure rate and coefficient of variation. This leads to an exaggerated magnitude of potentiation after optoPTP induction in FMRP DG-cKO mice.

### Presynaptic loss of FMRP induces alteration of Ca^2+^-related mechanisms at Mf synapses

Since Ca^2+^ elevations are slowly decaying in Mf terminals (Chamberland et al., 2018; Scott & Rusakov, 2006), in part due to elevated presynaptic Ca^2+^ buffering (Vyleta & Jonas, 2014), they constitute optimal conditions for asynchronous glutamate release (Chamberland et al., 2020).

Asynchronous release typically begins ∼10 ms after the action potential and can persist for several hundreds of milliseconds. Like synchronous release, it is Ca^2+^-dependent, but it specifically relies on the residual Ca^2+^ that accumulates in the presynaptic terminal after the action potential (Rozov et al., 2019). Thus, asynchronous release is a proxy for residual Ca^2+^ accumulation at the presynapse. In order to know whether presynaptic loss of FMRP impairs Ca^2+^ accumulation in Mf boutons we measured the asynchronous charge 20 ms after light stimulation during 0.1, 1, 3, and 20 Hz protocols (either 10 or 50 stimulations in trains) (**Figure 5A, C, E**). For 1 Hz and 3 Hz stimulation frequencies, asynchronous charge was reduced in FMRP DG-cKO mice compared to control mice (WT: *n*=15, cKO: *n*=16; *F*(1, 29)=6.7, *p*=0.0152; **Figure 5B**). The asynchronous charge after short trains was also reduced in FMRP DG-cKO mice compared to control mice (WT: 8.0 ± 4.4, *n*=15; cKO: 4.7 ± 2.6, *n*=16; *t*(22.2)=2.5, *p*=0.0183; **Figure 5D**) but asynchronous charge after optoPTP induction was no longer different between groups (WT: 17.9 ± 10.1, *n*=23; cKO: 18.7 ± 23, *n*=28; *U*=279, *p*=0.4244; **Figure 5F**). These results reveal that upon increase in frequency, residual Ca^2+^ accumulates less at the presynaptic terminal in the absence of presynaptic FMRP resulting in a decrease in Mf-EPSCs amplitude (**Figure 3B, G, H, L**). However, after optoPTP induction residual Ca^2+^ seems normalized at the same level in FMRP DG-cKO and control mice resulting in similar Mf-EPSCs amplitude right after induction (**Figure 4E**).

**Figure 5.**
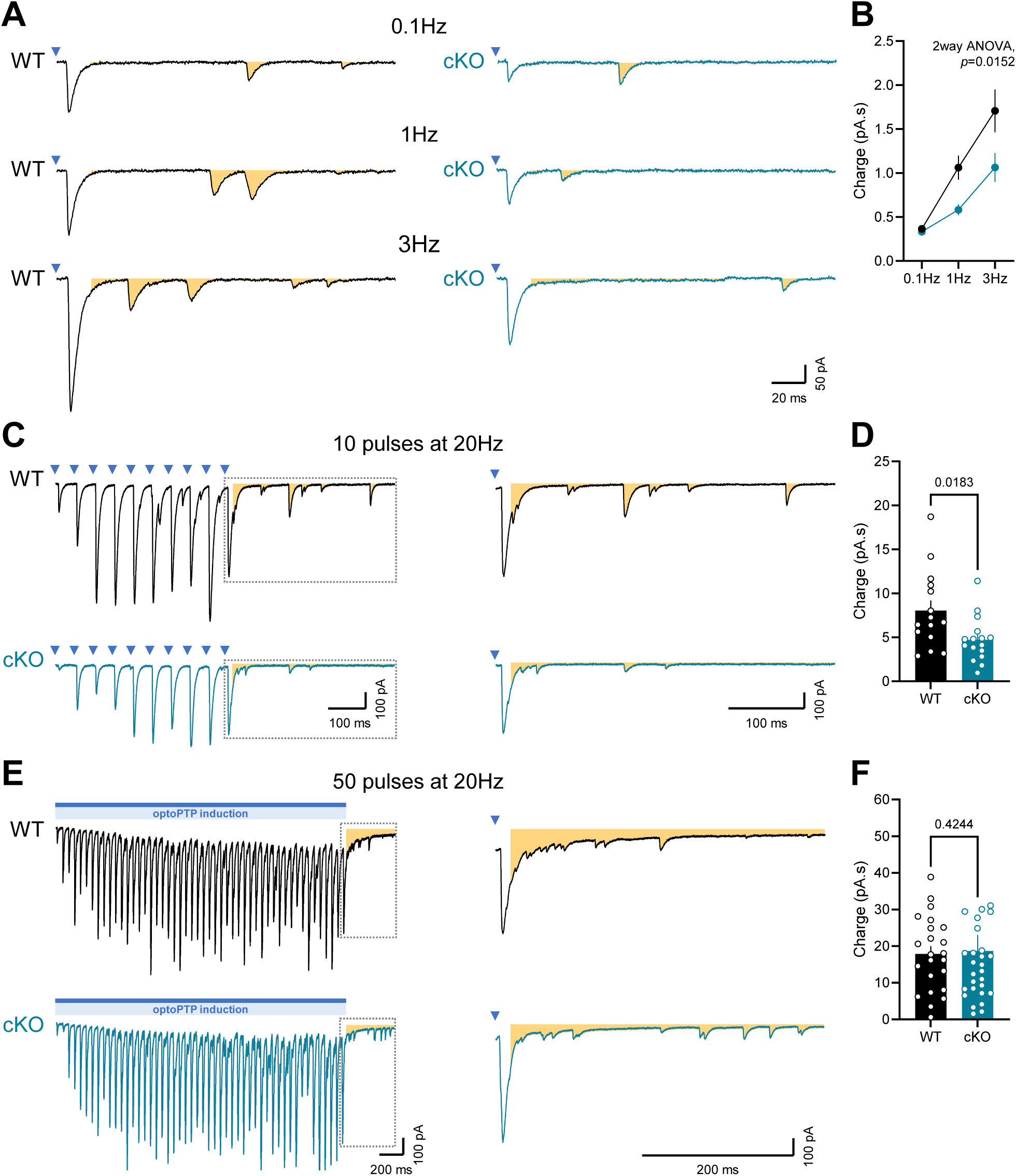
Reduction of asynchronous charge is abolished after optoPTP induction in the absence of presynaptic FMRP. (A) Representative 200 ms traces after light pulse, during protocols of 0.1 Hz, 1 Hz and 3 Hz. Asynchronous charge (orange area) has been calculated from 20 ms after light pulse. Recordings from CA3 pyramidal neurons in acute brain slices from control (black) and FMRP DG-cKO (blue) mice. (B) Mean asynchronous charge is decreased in the absence of presynaptic FMRP during 1 Hz and 3 Hz protocols (2way ANOVA, main effect between groups, *F*(1, 29)=6.7, *p*=0.0152). WT: *n*=15 cells; cKO: *n*=16 cells. WT: *N*=8 mice; cKO: *N*=10 mice. (C) *Left panels*: Representative traces of Mf-EPSCs obtained during a train of 10 light pulses at 20 Hz from CA3 pyramidal neurons in acute brain slices from control and FMRP DG-cKO mice. *Right panels*: Close-up of boxed areas showing 450 ms traces after the last light pulse. Asynchronous charge (orange area) has been calculated from 20 ms after light pulse. (D) Mean asynchronous charge is decreased in the absence of presynaptic FMRP after a train of 10 light pulses at 20 Hz (Welch’s t-test, *t*(22.2)=2.5, *p*=0.0183). WT: *n*=15 cells; cKO: *n*=16 cells. WT: *N*=8 mice; cKO: *N*=10 mice. (E) *Left panels*: Representative traces of Mf-EPSCs obtained during a train of 50 light pulses at 20 Hz from CA3 pyramidal neurons in acute brain slices from control and FMRP DG-cKO mice. *Right panels*: Close-up of boxed areas showing 450 ms traces after the last light pulse. Asynchronous charge (orange area) has been calculated from 20 ms after light pulse. (F) Mean asynchronous charge is not altered in the absence of presynaptic FMRP after a train of 50 light pulses at 20Hz (Mann-Whitney test, *U*=279, *p*=0.4244). WT: *n*=23 cells; cKO: *n*=28 cells. WT: *N*=14 mice; cKO: *N*=20 mice. Data are shown as mean ± SEM.

**Figure 6.**
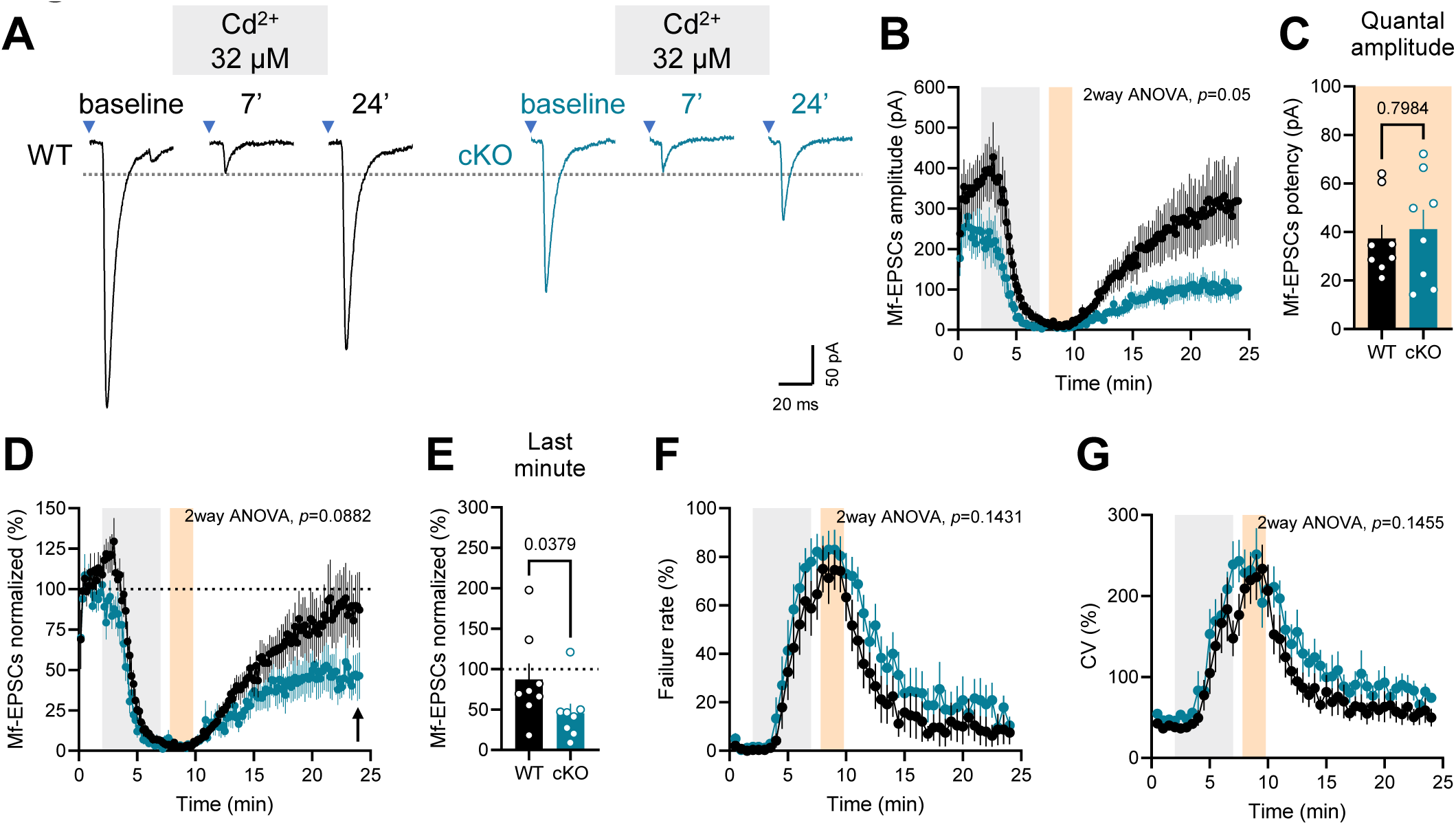
Quantal amplitude is unaltered, but recovery from VGCCs inhibition is impaired in the absence of presynaptic FMRP. (A) Representative traces of Mf-EPSCs recorded at 1 Hz during 2 min baseline, after 5 min application of cadmium (32 µM, voltage-gated calcium channels inhibitor, grey area), and 17 min after cadmium application (washout). Recordings from CA3 pyramidal neurons in acute brain slices from control (black) and FMRP DG-cKO (blue) mice. Dashed line indicates average quantal amplitude. (B) Time course of Mf-EPSCs amplitude for the experiment in (A) shows decreased amplitudes in the absence of presynaptic FMRP (2way ANOVA, main effect between groups, *F*(1, 14)=4.5, *p*=0.05). Time points are made by doing the average amplitude of 10 Mf-EPSCs. The orange area shows where the quantal amplitude analysis has been performed (see Methods). (C) Histogram showing similar quantal amplitudes (Mann-Whitney test, *U*=29, *p*=0.7984) calculated within the orange area and without considering failures. (D) Time course of normalized Mf-EPSCs amplitude for the experiment in (A) shows not statistically different amplitudes in the absence of presynaptic FMRP (2way ANOVA, main effect between groups, *F*(1, 14)=3.4, *p*=0.0882). Time points are made by doing the average amplitude of 10 Mf-EPSCs. (E) Histogram of normalized Mf-EPSCs amplitude calculated during the last minute of recording (arrow, D) showing decreased recovery from cadmium application in the absence of presynaptic FMRP (Mann-Whitney test, *U*=12, *p*=0.0379). (F) Time course of failure rate of Mf-EPSCs for the experiment in (A) is not statistically different between WT and cKO (2way ANOVA, main effect between groups, *F*(1, 14)=2.4, *p*=0.1431). Time points are made by calculating failure rate during 30 sec of recording. (G) Time course of coefficient of variation of Mf-EPSCs for the experiment in (A) is not statistically different between control and FMRP DG-cKO (2way ANOVA, main effect between groups, *F*(1, 14)=2.4, *p*=0.1455). Time points are made by calculating coefficient of variation during 30 sec of recording. WT: *n*=8 cells; cKO: *n*=8 cells. WT: *N*=4 mice; cKO: *N*=4 mice. Data are shown as mean ± SEM.

Another finding establishes a link between presynaptic FMRP and Ca^2+^-related mechanisms: recovery following inhibition of VGCCs. After adding Cd^2+^ to the bath, we recorded the washout period to ensure complete recovery of the Mf-EPSCs amplitude following Ca^2+^ channel inhibition by Cd^2+^ (**Figure 6A**). We expected both groups to recover completely. However, the normalized Mf-EPSCs relative amplitude was different between groups (WT: 87.3 ± 55.2, *n*=8; cKO: 45.2 ± 33.9, *n*=8; *U*=12, *p*=0.0379; **Figure 6E**). Further statistical analyses showed that normalized Mf-EPSCs amplitudes in control mice fully recovered at the end of the experiment (*T*=-10, *p*=0.5469), whereas normalized Mf-EPSCs amplitudes in FMRP DG-cKO mice only partially recovered (*T*=-34, *p*=0.0156). Altogether these results indicate that loss of presynaptic FMRP impairs Ca^2+^ accumulation and dynamics in Mf terminals.

### Presynaptic loss of FMRP deregulates AC-cAMP-PKA signaling pathway at Mf synapses

Ca^2+^ influx can stimulate cAMP production through Ca^2+^/calmodulin-activated adenylyl cyclases (ACs), leading to activation of protein kinase A (PKA), a key regulator of synaptic plasticity (Ferguson & Storm, 2004; H. Wang & Storm, 2003; Wong et al., 1999). At hippocampal Mf terminals, PKA is required for the induction and maintenance of PTP (Y. Y. Huang et al., 1994; Kim et al., 2024; Vandael et al., 2020; Weisskopf et al., 1994). Because optoPTP induction was the only protocol that rescued the presynaptic phenotype observed in FMRP DG-cKO mice, we next asked whether PKA activity also contributes to basal neurotransmission in the absence of presynaptic FMRP.

After a baseline of 10 minutes at 0.1 Hz, Mf-EPSCs were recorded for 50 minutes in the presence of the PKA inhibitor H89, at a concentration of 10 µM (**Figure 7A**). In control mice, H89 application caused a marked reduction of Mf-EPSCs amplitude. After 45 minutes of recording, amplitude had declined to a level comparable to the baseline amplitude of Mf-EPSCs from FMRP DG-cKO mice (blue dashed line; **Figure 7B**). Across the entire recording, absolute Mf-EPSCs amplitude was lower in FMRP DG-cKO than in control mice (WT: *n*=13, cKO: *n*=16; *F*(1, 27)=13.5, *p*=0.001; **Figure 7B**). It is important to note that the effect of H89 observed in control mice using optogenetic stimulation was replicated using electrical stimulation (**Figure S2C-F**). When normalized to baseline, however, Mf-EPSCs relative amplitude was significantly higher in FMRP DG-cKO compared to control mice (WT: *n*=13, cKO: *n*=16; *F*(1, 27)=4.9, *p*=0.0358; **Figure 7C**). Notably, after 30 minutes of H89 application, Mf-EPSCs from FMRP DG-cKO mice were still at ∼93% of baseline amplitude, whereas Mf-EPSCs from control mice declined to ∼65% of baseline (WT: 65.2 ± 14.6, *n*=13; cKO: 92.6 ± 30.4, *n*=16; Šídák’s multiple comparisons test*p*=0.0254). Following H89 application, the failure rate (WT: *n*=13, cKO: *n*=16; *F*(1, 27)=1.5, *p*=0.2286; **Figure 7D**) and the coefficient of variation (WT: *n*=13, cKO: *n*=16; *F*(1, 27)=1.5, *p*=0.2246; **Figure 7E**) did not differ significantly between genotypes. The PPRs were not different between FMRP DG-cKO and control mice following H89 application (**Figure S2A, B**).

**Figure 7.**
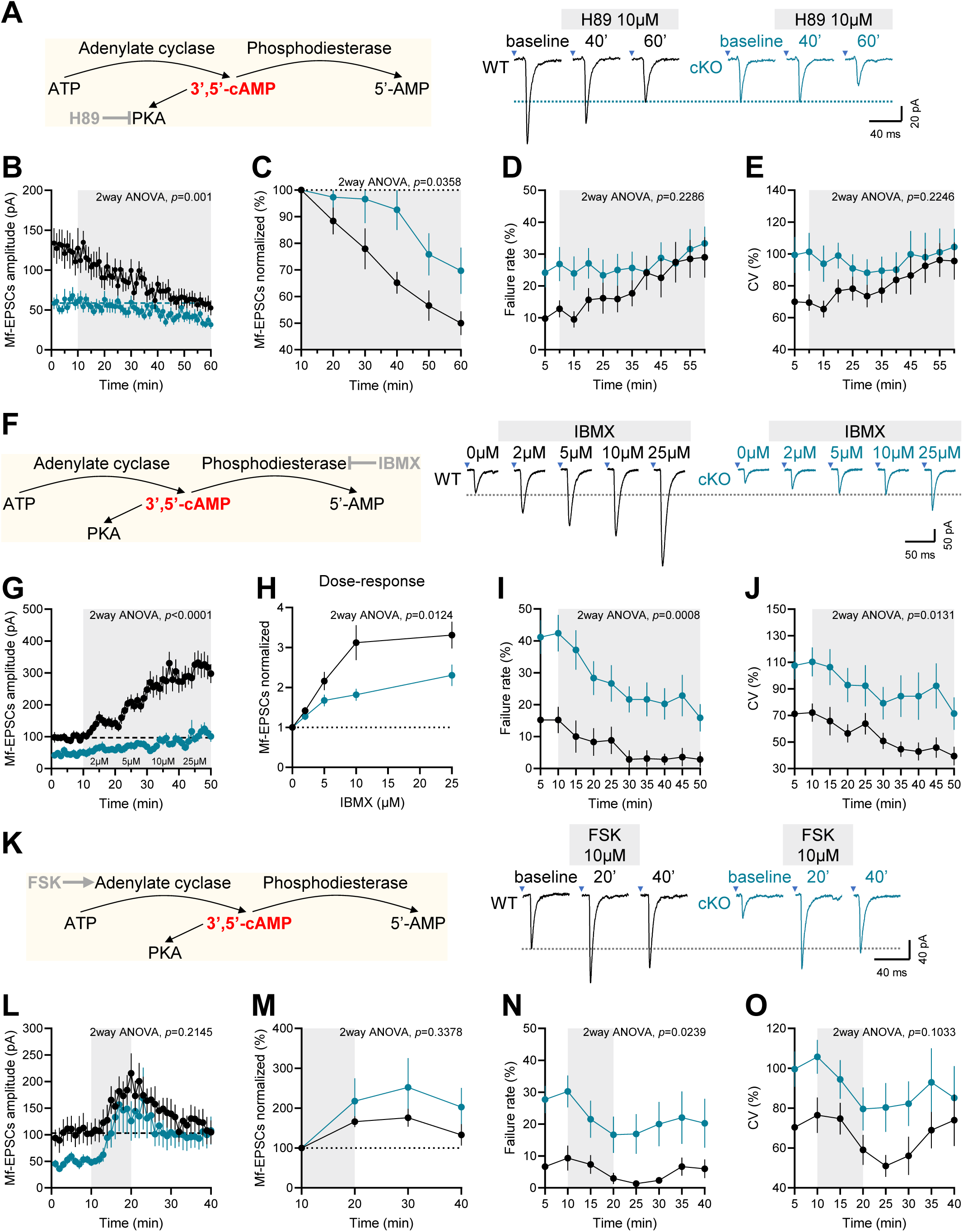
AC-cAMP-PKA pathway is impaired during basal synaptic transmission in the absence of presynaptic FMRP. (A) *Left panel*: cAMP-related signaling pathway. Adenylate cyclase (AC) catalyzes the conversion of adenosine triphosphate (ATP) to 3’,5’-cyclic AMP (cAMP). Then, cAMP activates various effectors such as the cAMP-dependent protein kinase (PKA). Finally, phosphodiesterase (PDE) degrades the phosphodiester bond in the cAMP to produce 5’-AMP. H89 is an inhibitor of PKA. *Right panel*: Representative traces of Mf-EPSCs recorded at 0.1 Hz during 10 min baseline, 30 min and 50 min after H89 application (10 µM, grey area). Recordings from CA3 pyramidal neurons in acute brain slices from control (black) and FMRP DG-cKO (blue) mice. Blue dashed line indicates average baseline amplitude obtained from FMRP DG-cKO mice. (B) Time course of Mf-EPSCs amplitude for the experiment in (A) shows decreased amplitudes in the absence of presynaptic FMRP (2way ANOVA, main effect between groups, *F*(1, 27)=13.5, *p*=0.001). Time points are made by doing the average amplitude of 6 Mf-EPSCs. Blue dashed line indicates average baseline amplitude obtained from FMRP DG-cKO mice. (C) Time course of normalized Mf-EPSCs amplitude for the experiment in (A) shows decreased amplitudes in control mice during H89 application (2way ANOVA, main effect between groups, *F*(1, 27)=4.9, *p*=0.0358). Time points are made by doing the average normalized amplitude of Mf-EPSCs during 10 min. (D) Time course of failure rate of Mf-EPSCs for the experiment in (A) is not statistically different between control and FMRP DG-cKO mice during H89 application (2way ANOVA, main effect between groups, *F*(1, 27)=1.5, *p*=0.2286). Time points are made by calculating failure rate during 5 min of recording. (E) Time course of coefficient of variation of Mf-EPSCs for the experiment in (A) is not statistically different between control and FMRP DG-cKO mice during H89 application (2way ANOVA, main effect between groups, *F*(1, 27)=1.5, *p*=0.2246). Time points are made by calculating coefficient of variation during 5 min of recording. WT: *n*=13 cells; cKO: *n*=16 cells. WT: *N*=11 mice; cKO: *N*=15 mice. (F) *Left panel*: cAMP-related signaling pathway. AC catalyzes the conversion of ATP to 3’,5’-cAMP. Then, cAMP activates various effectors such as PKA. Finally, PDE degrades the phosphodiester bond in the cAMP to produce 5’-AMP. IBMX is an inhibitor of PDEs. *Right panel*: Representative traces of Mf-EPSCs recorded at 0.1 Hz during 10 min baseline, after 10 min application of 2, 5, 10 and 25 µM of IBMX (grey area). Recordings from CA3 pyramidal neurons in acute brain slices from control and FMRP DG-cKO mice. Dashed line indicates average baseline amplitude of control mice. (G) Time course of Mf-EPSCs amplitude for the experiment in (F) shows decreased amplitudes in the absence of presynaptic FMRP (2way ANOVA, main effect between groups, *F*(1, 26)=44, *p*=0.001). Time points are made by doing the average amplitude of 6 Mf-EPSCs. (H) Summary of the dose response experiment in (F) of normalized Mf-EPSCs amplitudes as a function of IBMX concentration showing decreased amplitudes in the absence of presynaptic FMRP (2way ANOVA, main effect between groups, *F*(1, 26)=7.2, *p*=0.0124). (I) Time course of failure rate of Mf-EPSCs for the experiment in (F) shows a sustained increase during IBMX application (2way ANOVA, main effect between groups, *F*(1, 26)=14.3, *p*=0.0008). Time points are made by calculating failure rate during 5 min of recording. (J) Time course of coefficient of variation of Mf-EPSCs for the experiment in (F) shows a sustained increase during IBMX application (2way ANOVA, main effect between groups, *F*(1, 26)=7, *p*=0.0131). Time points are made by calculating coefficient of variation during 5 min of recording. WT: *n*=14 cells; cKO: *n*=14 cells. WT: *N*=10 mice; cKO: *N*=10 mice. (K) *Left panel*: cAMP-related signaling pathway. AC catalyzes the conversion of ATP to 3’,5’-cAMP. Then, cAMP activates various effectors such as PKA. Finally, PDE degrades the phosphodiester bond in the cAMP to produce 5’-AMP. Forskolin (FSK) is an activator of ACs. *Right panel*: Representative traces of Mf-EPSCs recorded at 0.1 Hz during 10 min baseline, after 10 min application of FSK (10 µM, grey area), and 20 min after FSK application (washout). Recordings from CA3 pyramidal neurons in acute brain slices from control and FMRP DG-cKO mice. Dashed line indicates average baseline amplitude of control mice. (L) Time course of Mf-EPSCs amplitude for the experiment in (K) shows similar amplitudes in the absence of presynaptic FMRP (2way ANOVA, main effect between groups, *F*(1, 21)=1.6, *p*=0.2145). Time points are made by doing the average amplitude of 6 Mf-EPSCs. (M) Time course of normalized Mf-EPSCs amplitude for the experiment in (K) shows similar amplitudes in the absence of presynaptic FMRP (2way ANOVA, main effect between groups, *F*(1, 21)=1, *p*=0.3378). Time points are made by doing the average normalized amplitude of Mf-EPSCs during 10 min. (N) Time course of failure rate of Mf-EPSCs for the experiment in (K) is increased in the absence of presynaptic FMRP (2way ANOVA, main effect between groups, *F*(1, 21)=6, *p*=0.0239). Time points are made by calculating failure rate during 5 min of recording. (O) Time course of coefficient of variation of Mf-EPSCs for the experiment in (K) is not statistically different between control and FMRP DG-cKO mice (2way ANOVA, main effect between groups, *F*(1, 21)=3, *p*=0.1033). Time points are made by calculating coefficient of variation during 5 min of recording. WT: *n*=10 cells; cKO: *n*=13 cells. WT: *N*=8 mice; cKO: *N*=10 mice. Data are shown as mean ± SEM.

Overall, these experiments indicate that inhibition of PKA mimicked the FMRP DG-cKO phenotype in control mice, reducing Mf-EPSCs amplitude together with increasing the failure rate and coefficient of variation. This suggests that PKA is very likely less active in FMRP DG-cKO mice during basal synaptic activity. This effect can be explained by three hypotheses: (1) PKA is less active due to more degradation of cAMP mediated by phosphodiesterases (PDEs), or (2) due to less activity of ACs, or (3) both mechanisms are conjugated to impair PKA activity. The first hypothesis has been tested using IBMX a non-selective PDEs inhibitor. After a baseline of 10 minutes at 0.1 Hz, Mf-EPSCs were recorded for 40 minutes in the presence of IBMX at different successive concentrations: 2, 5, 10 and 25 µM (**Figure 7F**). In control mice, IBMX application caused a marked enhancement of Mf-EPSCs amplitude. However, across the entire recording, absolute Mf-EPSCs amplitude was lower in FMRP DG-cKO than in control mice (WT: *n*=14, cKO: *n*=14; *F*(1, 26)=44, *p*<0.0001; **Figure 7G**). When normalized to baseline, Mf-EPSCs relative amplitude was still significantly higher in control compared to FMRP DG-cKO mice (WT: *n*=14, cKO: *n*=14; *F*(1, 26)=7.2, *p*=0.0124; **Figure 7H**). Finally, IBMX application did not normalize the failure rate (WT: *n*=14, cKO: *n*=14; *F*(1, 26)=14.3, *p*=0.0008; **Figure 7I**) nor the coefficient of variation (WT: *n*=14, cKO: *n*=14; *F*(1, 26)=7, *p*=0.0131; **Figure 7J**) deficits observed in FMRP DG-cKO mice. Therefore, an increased activity of PDEs is rather unlikely, because the phenotype is still present in FMRP DG-cKO mice upon IBMX application.

The role of ACs has been tested using forskolin, a non-selective ACs activator at a concentration of 10 µM. After a baseline of 10 minutes at 0.1 Hz, Mf-EPSCs were recorded for 10 minutes in the presence of FSK and then during 20 minutes of washout (**Figure 7K**). In both control and and FMRP-cKO mice, FSK application caused a marked enhancement of Mf-EPSCs amplitude. After FSK application, absolute Mf-EPSCs amplitude was not statistically different between FMRP DG-cKO and control mice (WT: *n*=10, cKO: *n*=13; *F*(1, 21)=1.6, *p*=0.2145; **Figure 7L**). However, when normalized to baseline, Mf-EPSCs relative amplitude was not statistically different in control compared to FMRP DG-cKO mice (WT: *n*=10, cKO: *n*=13; *F*(1, 21)=1, *p*=0.3378; **Figure 7M**). Finally, FSK application rescued the coefficient of variation (WT: *n*=10, cKO: *n*=13; *F*(1, 21)=3, *p*=0.1033; **Figure 7O**) but not the failure rate (WT: *n*=10, cKO: *n*=13; *F*(1, 21)=6, *p*=0.0239; **Figure 7N**) deficit observed in FMRP DG-cKO mice. In conclusion, these results suggest that presynaptic FMRP impairs presynaptic ACs activity, because the phenotype is almost completely rescued in FMRP DG-cKO mice upon FSK application.

## Discussion

In this study, we demonstrate that selective presynaptic deletion of FMRP at hippocampal Mf synapses impairs basal neurotransmission, as evidenced by reduced Mf-EPSC amplitude, increased failure rate and altered coefficient of variation, through presynaptic mechanisms. Our data further implicate the cAMP-PKA signaling pathway as a potential cause of this phenotype, likely leading to a reduction in the RRP size at AZs.

Unexpectedly, the conditional deletion of FMRP did not lead to altered intrinsic excitability of DG cells, contrasting with the hyperexcitability reported in constitutive FMRP-deficient models (Contractor et al., 2015; Deng & Klyachko, 2021; Liu et al., 2022). This difference may result from the timing of FMRP deletion (Gredell et al., 2023). Whereas previous studies used constitutive *Fmr1*-KO or mosaic models, we deleted FMRP after maturation of DG cells and their Mf synapses (Amaral & Dent, 1981; Marchal & Mulle, 2004), thereby minimizing developmental adaptations. Our results may therefore better reflect the physiological functions of presynaptic FMRP rather than secondary consequences of its developmental absence. Constitutive models necessarily combine the immediate effects of FMRP loss with long-term compensatory mechanisms arising during circuit formation and maturation. In contrast, post-developmental deletion provides a means of isolating primary physiological functions of FMRP. This distinction may explain discrepancies with the classical Fragile X literature and suggests that some phenotypes associated with Fragile X syndrome could result from the absence of FMRP during development in addition to its direct loss at the adult stage.

A consistent finding across the study was the marked (∼50%) reduction in Mf-EPSC amplitude during basal synaptic transmission, together with unaltered short-term facilitation, in mice lacking presynaptic FMRP. These findings extend previous observations at an excitatory-to-inhibitory synapse in the mouse neocortex (Patel et al., 2013) to an excitatory-to-excitatory hippocampal synapse. Several observations indicate that the defect is presynaptic: amplitude covaried with the coefficient of variation of Mf-EPSCs, whereas PPR and quantal amplitude were unchanged. The increased failure rate, together with an identical number of bassoon-containing active zones between groups, further suggests a reduction in the number of release-ready vesicles per AZ and therefore in RRP size in the absence of presynaptic FMRP.

Because PKA has been implicated in synaptic vesicle docking and priming and in regulating RRP size at AZs (Bruentgens et al., 2026; Cazares et al., 2016; Cesca et al., 2010; Leenders & Sheng, 2005; Pernía-Andrade & Jonas, 2011), particularly during PTP at Mf synapses (Kim et al., 2024; Vandael et al., 2020), we investigated whether presynaptic FMRP loss alters this form of plasticity. OptoPTP magnitude was increased in the absence of presynaptic FMRP, yet Mf-EPSCs amplitude immediately after optoPTP induction was similar between groups. The asynchronous charge following induction was normalized in FMRP DG-cKO mice, suggesting comparable Ca^2+^ accumulation in Mf terminals during the induction train. In contrast, asynchronous charge was reduced at 1 Hz, 3 Hz and after a 10-pulse train at 20 Hz in the absence of presynaptic FMRP.

The relationship between FMRP and presynaptic voltage-gated calcium channels (VGCCs) provides a potential mechanistic explanation. FMRP regulates surface expression and conductance of Cav2 channels: its loss decreases Cav2.1 (P/Q-type) surface expression and conductance (Castagnola et al., 2018), while increasing Cav2.2 (N-type) and Cav2.3 (R-type) expression and conductance (Ferron et al., 2014, 2020; Gray et al., 2019). At Mf boutons, these isoforms contribute differently depending on activity. During a single action potential and short trains (<12 APs), most Ca^2+^ influx is mediated by Cav2.1, followed by Cav2.2 and a smaller Cav2.3 contribution (Breustedt et al., 2003; L. Li et al., 2007). As stimulation increases, the contribution of Cav2.3 progressively rises, becoming dominant from approximately 20 APs and supporting PTP induction (Dietrich et al., 2003; Vandael et al., 2020).

Based on these observations, we propose that presynaptic FMRP regulates neurotransmitter release through activity-dependent control of Cav2 channels and cAMP-PKA signaling. During basal activity and short trains, reduced Cav2.1 expression in the absence of FMRP would decrease Ca^2+^ influx at Mf terminals (Castagnola et al., 2018; Chamberland et al., 2017, 2020; L. Li et al., 2007). This reduction would decrease activation of the Ca^2+^/CaM-dependent adenylyl cyclases AC1 and AC8 (Shahoha et al., 2022), thereby reducing PKA activity and consequently synaptic vesicle docking/priming and RRP size. During optoPTP induction, however, Ca^2+^ influx is predominantly mediated by Cav2.3, which is overexpressed in the absence of FMRP (Gray et al., 2019). Enhanced Cav2.3-dependent Ca^2+^ entry could therefore increase AC1/8 activation and PKA activity during the induction train, increasing the number of docked/primed vesicles and explaining the enhanced optoPTP magnitude in FMRP DG-cKO mice.

Pharmacological experiments support this framework. H89, a PKA inhibitor, reproduced the FMRP DG-cKO phenotype in control mice, whereas IBMX-mediated PDE inhibition did not reverse the phenotype in FMRP DG-cKO mice. In contrast, activation of ACs with forskolin almost completely restored basal synaptic activity. These results suggest that, in the absence of presynaptic FMRP, cAMP production by ACs and subsequent PKA recruitment are reduced during basal activity, resulting in impaired vesicle docking/priming. We also identify a previously underappreciated role for presynaptic PKA in maintaining efficient basal transmission at mature Mf synapses, which has generally been associated with high-frequency activity such as PTP and LTP (Vandael et al., 2020; Weisskopf et al., 1994).

We also identified impaired recovery of synaptic activity following VGCC inhibition in the absence of presynaptic FMRP. To our knowledge, this effect has not previously been reported following Cd^2+^ application. Reduced VGCC surface expression alone, particularly of Cav2.1, may not fully explain this phenotype and could instead reflect altered channel dynamics. One possibility involves changes in Cav2 auxiliary subunits: α2δ and β subunits regulate Cav2 trafficking and kinetics (Dolphin, 2013, 2016, 2018; Isom et al., 1994), and reduced expression could slow VGCC recovery after inhibition. Another possibility involves the clathrin-adaptor protein complex 2 (AP-2), whose expression is normally repressed by FMRP and which mediates endocytosis of synaptic proteins including the α2δ-1 VGCC subunit (Shi et al., 2025). Increased AP-2 expression in the absence of FMRP could enhance activity-dependent endocytosis of α2δ-1-associated VGCCs (Imoto & Watanabe, 2025; Kadlecova et al., 2016; McMahon & Boucrot, 2011; Royle & Lagnado, 2010). Because Cav2.1 is preferentially coupled to α2δ-1 whereas Cav2.3 is not (Dolphin, 2016), this mechanism could account for both the presynaptic phenotype and impaired recovery following Cd^2+^ application.

Several limitations should be considered. First, our results indirectly indicate that Cav2 channels are altered at Mf terminals, but direct confirmation of Cav2.1 surface loss and Cav2.3 overexpression, as well as measurements of channel-specific Ca^2+^ conductance, are required, ideally using whole-cell patch-clamp recordings from Mf boutons. Second, although our pharmacological results suggest reduced PKA activity during basal transmission, H89 can affect targets other than PKA (Y. Chen & Sabatini, 2021; Davies et al., 2000; Fusi et al., 2017; Limbutara et al., 2019; Lochner & Moolman, 2006; Murray, 2008). Other PKA inhibitors, such as Rp-8-CPT-cAMPS or Rp-8-Br-cAMPS, and cAMP analogs such as Sp-cAMPS should therefore be used for confirmation. Direct measurements of presynaptic cAMP and PKA dynamics using FRET-based biosensors, such as Epac-S^H187^, cAMPFIRE and AKARs, would further test this mechanism (Klarenbeek et al., 2015; Ma et al., 2018; Massengill et al., 2022). Third, although IBMX did not reveal a direct contribution of PDEs, it also affects adenosine receptors (Prince & Stevens, 1992); selective PDE subtype inhibitors will therefore be required to confirm this conclusion. Nevertheless, PDEs may still contribute under specific activity patterns, developmental stages, subcellular compartments or neuromodulatory conditions, consistent with reported links between FMRP and PDE2A, PDE4D and PDE10A. Fourth, the proposed reduction in RRP size should be directly tested by ultrastructural and functional approaches, including electron microscopy (e.g. cryo-ET, cryo-FIB/SEM, cryo-CLEM) to quantify docked vesicles and whole-cell recordings from Mf boutons to estimate RRP size from capacitance jumps (Bischofberger et al., 2006; Fukaya, Hirai, et al., 2023; Hallermann et al., 2003; Kushmerick & von Gersdorff, 2003; Midorikawa & Sakaba, 2017; Miyano et al., 2019).

Finally, an important question is whether the same presynaptic phenotype found in this study would occur after constitutive FMRP deletion. Replication in *Fmr1*-KO mice would support the conclusion that these mechanisms are a robust feature of presynaptic FMRP loss and directly contribute to Fragile X pathophysiology. Conversely, an absent or different phenotype would indicate that developmental compensation modifies the presynaptic consequences of FMRP deficiency. Identifying such mechanisms could clarify how synapses adapt to constitutive FMRP loss and reveal potential novel therapeutic targets.

Overall, our findings suggest that presynaptic FMRP integrates multiple signaling pathways converging on cAMP-PKA signaling to regulate neurotransmitter release during basal synaptic activity. Beyond their relevance to Fragile X syndrome, these findings provide a framework for understanding how presynaptic signaling pathways control vesicle release and synaptic function at mature Mf synapses. A deeper characterization of these mechanisms may therefore contribute to both fundamental neuroscience and the identification of new therapeutic strategies.

## Material and methods

### Animals

Male mice C57Bl/6J floxed for the *Fmr1* gene on the X chromosome (*Fmr1^flox/y^*), referred to as *Fmr1* conditional knockout (cKO) mice (Mientjes et al., 2006) and their non-floxed wild-type (WT) littermates were used in this study. Animals were generated from frozen sperm samples obtained from Janvier Labs (Le Genest-Saint-Isle, France) and generously provided by Prof. Isabelle Caillé† (Institute of Biology Paris-Seine, Sorbonne University, Paris, France). Breeding and housing were conducted at the EOPS and Pôle In Vivo-EXPE facilities respectively of the Centre Broca Nouvelle-Aquitaine (Bordeaux, France). All experimental procedures were approved by the local ‘Ethical Committee of Bordeaux’ (CEEA -050) and complied with European and French regulations on animal research (APAFIS no. 2021040211108206). Mice were weaned at postnatal day 21 (P21) and housed with their littermates, with *ad libitum* access to food and water. The cages were kept under controlled temperature conditions and a 12 hour light/dark cycle.

### Viral vector

Lentiviral vectors were used in this study because they enable efficient delivery of large transgenes into neurons with minimal cytotoxicity. After weaning (P25-P30), both floxed and non-floxed mice were stereotaxically injected in the dentate gyrus with the lentivirus LV.C1qL2.ChIEF-Tom-2A-Cre (Barthet et al., 2018). Sparse expression was driven by the *C1qL2* promoter, which is selectively enriched in DG cells (Dalley et al., 2008; Iijima et al., 2010). The construct allowed co-expression of ChIEF (a light-sensitive channelrhodopsin variant) and tdTomato in both WT and cKO mice, while Cre recombinase was separated from ChIEF and tdTomato by a self-cleaving 2A peptide sequence. The viral titer was approximately 1.21×10^9^ particles/mL.

### Stereotaxic injections

For analgesia, mice received a subcutaneous injection of lurocaine (7 mg/kg) at the incision site, followed by an intraperitoneal injection of buprenorphine (0.5 mg/kg) and meloxicam (5 mg/kg). Anesthesia was induced with 4% isoflurane and maintained at 1.5-2% (∼0.2 L/min), adjusted according to the animal’s respiratory rate. Mice were placed on a heating pad throughout the surgery to prevent hypothermia. After verifying the absence of pain reflexes, the incision site was shaved and then cleaned with povidone-iodine. Head alignment was adjusted using the anatomical bregma as reference.

A small craniotomy was performed at the following stereotaxic coordinates relative to bregma: AP = -2.2 mm, ML = ±2.0 mm, DV = -2.0 mm. 750 nL of lentivirus was injected per hemisphere at a rate of 100 nL/min using a microinjection pump (World Precision Instruments). Viral titers were optimized to maximize infection efficiency in the dentate gyrus. After injection, the incision was sutured and mice were allowed to recover in a heated cage under supervision. Body weight was monitored daily for at least 3 days or until full recovery to pre-surgery weight. Mice were used for electrophysiological recordings >1 month (minimum 35 and maximum 56 days) post-injection to ensure stable transgene expression.

### Immunohistochemistry

Mice were deeply anesthetized and transcardially perfused with 4% paraformaldehyde (PFA) prepared in 1× phosphate-buffered saline (PBS). Brains were carefully removed, post-fixed in 4% PFA for several hours or overnight, and sectioned coronally at a thickness of 50 μm using a vibratome (Leica Microsystems, VT1200S).

Free-floating sections were incubated for 1 hour at room temperature in a blocking solution containing 5% normal goat serum (NGS) and 0.3% Triton X-100 in PBS to reduce non-specific binding and permeabilize cell membranes. Sections were then incubated overnight at 4°C with primary antibodies diluted in blocking solution. The following primary antibodies were used: rabbit anti-FMRP (Rb11, home-made, 1:250) generously provided by Dr. Barbara Bardoni and Marielle Jarjat (Institut de pharmacologie moléculaire et cellulaire, Côte d’Azur University, Valbonne, France) and guinea pig anti-Cre recombinase (Synaptic Systems, 257004, 1:500).

The following day, sections were washed three times for 10 min in PBS and incubated for 2 hours at room temperature with secondary antibodies diluted in PBS. Secondary antibodies included goat anti-guinea pig Alexa Fluor 488 (1:500) and goat anti-rabbit Alexa Fluor 647 (1:500). Following three additional washes in PBS, sections were mounted onto glass slides, and coverslipped using Fluoromount mounting medium with DAPI (300 mM). Confocal images were acquired immediately after mounting.

### *Ex-vivo* patch-clamp recordings

Acute hippocampal slices were prepared from 2 month-old (P60-P86) mice previously injected with the lentiviral construct. Mice were deeply anesthetized by intraperitoneal injection of ketamine (100 mg/kg) and xylazine (20 mg/kg). After confirming the absence of pain reflexes, the animal was intracardially perfused through the left ventricle with ice-cold, oxygenated (95% O2, 5% CO2) artificial cerebrospinal fluid (ACSF) containing (in mM): 120 NaCl, 2.5 KCl, 1.25 NaH2PO4-H2O, 27 NaHCO3, 1 MgCl2 anhydrous, 2 CaCl2-H2O2, 16.5 glucose, 2.8 pyruvic acid, and 0.5 ascorbic acid (pH 7.4, 310 mOsm). The brain was rapidly removed, and the cerebellum and olfactory bulbs were cut out. The hemispheres were separated along the midline, and each medial side was glued onto the vibratome stage for parasagittal sectioning. Slices (300 µm) were cut in ice-cold, oxygenated ACSF using a vibratome (Leica Microsystems, VT1200S) and then incubated in ACSF at 34°C for 15 min recovery. Subsequently, slices were maintained at room temperature in continuously oxygenated ACSF for use within the same day.

Borosilicate glass pipettes (Harvard Apparatus) were pulled with a vertical puller (Narishige) to a resistance of 3-6 MΩ. For voltage-clamp recordings, electrodes were filled with a cesium-based internal solution containing (in mM): 125 Cs-methanesulfonate, 2 MgCl2-6H2O, 4 NaCl, 5 phosphocreatine, 10 EGTA, 10 HEPES, and 4 NaATP (pH 7.3 adjusted with CsOH, 300 mOsm).

For current-clamp recordings, a potassium-based internal solution was used (in mM): 115 K-gluconate, 10 KCl, 0.2 EGTA, 0.02 CaCl2, 10 HEPES, 15 phosphocreatine, 4 MgATP, and 0.3 NaGTP (pH 7.3 adjusted with KOH, 300 mOsm).

During voltage-clamp experiments, series resistance was monitored using a -10 mV, 5 ms test pulse at the start of each sweep. Recordings were discarded if series resistance exceeded 20 MΩ or varied by more than 20% relative to baseline for short recordings (i.e. less than 30 min), and 25 MΩ or varied by more than 30% relative to baseline for long recordings (i.e. more than 30 min). No correction for liquid junction potential nor series resistance compensation was applied. Neurons were kept at a holding potential of -70mV.

Signals were recorded using an EPC-10 amplifier (HEKA Elektronik), filtered at 2.9 kHz (Bessel), and digitized at 10 kHz. All recordings were performed at room temperature.

The hippocampus was first identified under 10× magnification (Nikon, Plan fluor 10×/0.30), and viral infection was verified by detecting tdTomato fluorescence. At higher magnification (Nikon, NIR Apo 60×/1.0), patch-clamp recordings were made. In voltage-clamp experiments CA3 PCs were patched, while optogenetic stimulation sites were localized in the dendritic tree of granule cells by real-time monitoring of evoked responses. The LED light intensity (470 nm) has been set at 66% (CoolLED, pE-1&2) because it reliably elicited action potentials in the soma of granule cells. To measure ‘light intensity’, we used the irradiance of the LED light at the focal plane (*Measuring Illumination Intensity with Accuracy and Precision*, 2019) in the same conditions as during optogenetic stimulation (i.e. in the presence of filters and dichroic mirror). Light power through the 60x objective has been measured with a power sensor (Thorlabs, S120C) and power meter (Thorlabs, PM100D). Light irradiance (E) in our conditions is then given by the following formula:

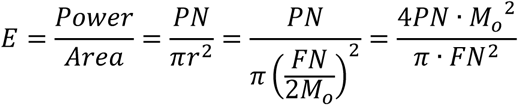

With the power number (PN, mW), field number (FN, mm) and objective magnification (Mo). For the objective used: FN is 22 mm and Mo is 60. At 66% of LED light intensity PN is 1.2 mW. Therefore, irradiance can be estimated around 11.36 mW/mm² at the focal plane during optogenetic stimulation in our experimental setup.

When we electrically stimulated mossy fiber tracts, we used minimal intensity stimulation via a borosilicate glass pipettes (∼1 µm tip diameter) positioned in the hilus region of the dentate gyrus. Stimulation pipettes were filled with ACSF and stimulation pulses in basal conditions were provided at 0.1 Hz (200 µs, 10-80 µA).

Monosynaptic activation of mossy fiber inputs was verified by the characteristic kinetics of the EPSCs and visual inspection of the waveform. Mf synaptic currents were identified according to the following criteria: robust low-frequency facilitation, low release probability at 0.1 Hz and rapid rise times of individual EPSCs. In current-clamp experiments DG granule cells were patched. Acquisition protocols were made and controlled using Patchmaster software (HEKA Elektronik).

The maximum number of cells recorded from a single animal was set at 5 to minimize statistical bias. When applying 20 Hz trains stimulation or pharmacological agents, only one cell was recorded per acute brain slice; in all other cases, the maximum number of cells recorded per slice was set at 2.

### Pharmacology

For voltage-clamp recordings, bicuculline (10 µM, Hellobio), CGP-55845 hydrochloride (1 µM, Hellobio), and D-APV (25 µM, Hellobio) were always included to ACSF in order to isolate AMPAR-and KAR-mediated conductance of the EPSCs.

All additional pharmacological agents were bath-applied in 100-200 mL of oxygenated ACSF using a closed-circuit system. The circuit was temporarily opened during the first 5 minutes of drug application to ensure the correct concentration reached the recording chamber, and similarly for the first 5 minutes of the washout period when applicable.

Additional pharmacological agents used in this study included forskolin (FSK, 10 µM, Sigma-Aldrich), H-89 dihydrochloride (10 µM, Tocris), 3-isobutyl-1-methylxanthine (IBMX, 2-25 µM, Sigma-Aldrich), and cadmium chloride (Cd^2+^, 32 µM, Sigma-Aldrich).

### Analysis of electrophysiological data

All analyses were performed using the Neuromatic plugin (Rothman & Silver, 2018) in IgorPro software (WaveMetrics). For voltage-clamp experiments on CA3 pyramidal cells, the amplitude, potency, failure rate and coefficient of variation of evoked excitatory postsynaptic currents (EPSCs) were measured. Amplitude of EPSCs (pA) is measured between the baseline current and the peak current, and its average calculation includes neurotransmission “failures” (i.e. 0 pA) if any. Potency of EPSCs (pA) is the average amplitude of EPSCs calculated without the “failures” (i.e. 0 pA). Failure rate (%) is the number of failures divided by the total number of EPSCs. Coefficient of variation (CV, %) is the standard deviation of EPSCs amplitude (σ) divided by the average of EPSCs amplitude (µ), as follows:

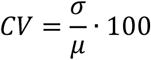

In order to study quantal amplitude at mossy fiber synapses we used 32 µM of Cd^2+^ (a calcium channels inhibitor) bath applied during a 1 Hz protocol of stimulation (Barthet et al., 2018). The rationale of this method is to drastically reduce the probability of release, so that the vast majority of recorded postsynaptic events result from mono-quantal release (∼95% of events). We recorded 2 minutes of baseline, then 5 minutes of Cd^2+^ application and finally 17 minutes of washout. We used the time window of maximum failure rate and CV to calculate the mean amplitude of successes (i.e. potency) in order to find mean quantal amplitude. The window was 2 minutes from 7′50″ to 9′50″ of the recording.

We quantified asynchronous charge (pA.s) following single action potential stimulation (0.1, 1, or 3 Hz) and trains of action potentials (10 or 50 pulses at 20 Hz). Asynchronous charge was used as an indirect measure of residual presynaptic calcium rather than to study asynchronous release itself (Llinares et al., 2025). Because it integrates the amplitude, frequency, and kinetics of asynchronous events, charge provides a global estimate of residual calcium-dependent release. For single action potentials, asynchronous charge was quantified over a 180 ms window beginning 20 ms after the light pulse. For action potential trains, asynchronous charge was quantified over a 430 ms window following the last stimulus beginning 20 ms after the light pulse. The initial 20 ms exclusion period after stimulation was applied to avoid contamination by synchronous release and was considered sufficient because, despite differences in evoked Mf-EPSCs amplitude between control and FMRP DG-cKO mice at 0.1 Hz (**Figure 2C**), the measured asynchronous charge was comparable between groups (**Figure 5B**).

For current-clamp experiments on DG granule cells, firing was evoked either by current steps or light pulses (1 ms at 470 nm). Infected WT or FMRP-cKO DG cells were patched as well as surrounding non-infected cells (NI). NI cells were patched regardless of their position inside the granule cell layer (i.e. outer or inner part). For current injection, 500 ms steps of hyperpolarizing currents (from -80 to 0 pA) and depolarizing currents (from 0 to 240 pA) were applied to cells with an increment of 20 pA. These steps were preceded by a 300 ms hyperpolarizing step of -20 pA. The average of membrane potentials recorded throughout the protocol during the steady-state of the 300 ms steps of -20 pA was used to calculate input resistance (MΩ). Resting membrane potential (RMP, mV) was calculated before the -20 pA step of the very first step recorded of the protocol (right after break-in). Rheobase (pA) was found from the first step of current eliciting at least one action potential, then the value of this parameter is the value of the current injected at that particular step. Spike rate (spikes/sec) was calculated by dividing the number of action potentials during a depolarizing current step by the duration of that step (here, 0.5 seconds). The action potential properties were analyzed on the first action potential elicited at the rheobase. Firing threshold (mV) was found using the first derivative of the action potential. Amplitude (mV) was calculated as the difference between the firing threshold to the peak. Full-width at half maximum (FWHM, ms) was measured as the time difference (Δt) between two points equal to half of the amplitude. Fast afterhyperpolarization amplitude (AHP, mV) was calculated as the difference between the firing threshold and the minimal value of potential after the repolarization phase (also known as AHP peak). For light stimulation, afterdepolarization amplitude (ADP, mV) was calculated as the difference between the AHP peak and the maximal value of potential after AHP peak (also known as ADP peak).

All analyses were performed blind to genotype and experimental group.

### STED and confocal imaging of Mf boutons

#### Tissue preparation and immunohistochemistry

Coronal hippocampal sections (70 μm) were prepared using a vibratome (VT1000S, Leica Microsystems) in 1× phosphate-buffered saline (PBS). Following dissection, the cerebellum and brainstem were removed, and brains were mounted on the vibratome stage for sectioning. Slices were collected in PBS and stored at 4°C until further processing.

For immunohistochemical labeling, free-floating sections were incubated for 2 hours at room temperature in a blocking solution containing 5% normal goat serum and 0.3% Triton X-100 in 1× PBS. Sections were subsequently washed three times in PBS and incubated overnight at 4°C with a mouse monoclonal anti-Bassoon antibody (Synaptic Systems, 141111) diluted 1:500 in blocking solution. The following day, sections were washed three times in PBS and incubated for 2 hours at room temperature with an ATTO647N-conjugated anti-mouse secondary antibody (Sigma-Aldrich, 50185) diluted 1:500 in blocking solution. Following additional washes in PBS, sections were mounted in Mowiol mounting medium for stimulated emission depletion (STED) microscopy.

#### STED microscopy

STED imaging was performed at the Bordeaux Imaging Centre (BIC, Bordeaux, France) using a Leica DMI6000 TCS SP8 X microscope. Images were acquired using Hybrid detectors with a scan speed of 400, line averaging set to four repetitions, a resolution of 1024 × 1024 pixels at 12-bit depth, and a zoom factor of 6, yielding a pixel size of 20 × 20 nm. Z-stacks encompassing the entire mossy fiber bouton were acquired for all samples.

TdTomato fluorescence was excited with a 561 nm laser and collected between 570 and 621 nm. Bassoon immunoreactivity was detected using excitation at 647 nm and emission collection between 670 and 720 nm. Super-resolution imaging was achieved using a 775 nm depletion laser.

#### Confocal microscopy

Confocal imaging was performed at the BIC using a Leica DM6 CFS TCS SP8 microscope equipped with a 63× oil-immersion objective. TdTomato fluorescence was excited at 561 nm and detected between 570 and 621 nm using photomultiplier tube (PMT) detectors. Acquisition parameters were maintained constant across all samples, including a scan speed of 400, a resolution of 1024 × 1024 pixels at 16-bit depth, and collection of Z-stacks spanning the entire volume of the tissue section.

#### Mf bouton volume analysis

Morphological analyses were performed using Imaris software (Oxford Instruments). Mossy fiber bouton volume was quantified using the Surface tool after defining an appropriate region of interest. Segmentation thresholds were adjusted according to infection density but maintained within a consistent range across samples. Objects with volumes smaller than 5 μm³ or larger than 70 μm³ were excluded from analysis. All analyses were performed blind to genotype and experimental group.

#### Bassoon clusters analysis

Bassoon clusters were quantified using Imaris software. Mossy fiber boutons were first segmented using the Surface tool, and bassoon-positive clusters located within the defined bouton volume were identified using the Spots tool. Detection parameters were adjusted according to staining intensity while remaining within a consistent range across all samples. Z-axis elongation resulting from stack acquisition was taken into account during clusters detection and quantification. Only the clusters found within 200 nm from the plasma membrane were counted. All analyses were performed blind to genotype and experimental group.

### Statistical analysis

Recordings of 0.1 Hz baselines, paired-pulses, 20 Hz trains, and optoPTP induction trains recorded in exact same experimental conditions at different time points during the whole investigation, were pooled together (when applicable) to perform statistical analyses and graph making.

Normality of data distributions was assessed using the Shapiro-Wilk test, which is well suited for the sample sizes used in this study (Ghasemi & Zahediasl, 2012). Depending on the outcome of the normality test, either parametric or non-parametric statistical tests were applied as appropriate. To compare two independent means, two-sided unpaired Welch’s t-test (*t*) or Mann-Whitney (*U*) tests were used. To compare cumulative distributions of two independent data sets, Kolmogorov-Smirnov (*D*) test was used. To compare individual means with a hypothetical mean, one-sample t-test (*t*) or Wilcoxon signed rank test (*T*) were used. To compare more than two independent means, Welch’s one-way ANOVA (*W*, followed by Dunnett’s post-hoc test) or Kruskal-Wallis (*H*, followed by Dunn’s T3 post-hoc test) tests were used. Two compare two or more independent means across multiple time points, 2way ANOVA (mixed model with Geisser-Greenhouse correction) test (*F*) was used, main effect between groups was reported, and when applicable this test was followed by Šídák’s multiple comparisons test or Fisher’s LSD post-hoc test. Statistical significance was set at *p≤*0.05. No outlier tests have been used. No statistical test was used to predetermine the sample size. All analyses and graph making were performed using GraphPad Prism (GraphPad Software).

Data are presented as mean ± SEM in graphical figures and as mean ± SD in the text of the ‘Results’ section.

## Data availability

All data generated in this study will be publicly available.

## Acknowledgments

We thank Prof. Isabelle Caillé† (Institute of Biology Paris-Seine, Sorbonne University, Paris, France) for providing sperm samples used to generate our mice. We thank Dr. Barbara Bardoni (Institut de pharmacologie moléculaire et cellulaire, Côte d’Azur University, Valbonne, France) for the insightfull discussions, and Marielle Jarjat for providing the anti-FMRP primary antibody used in the study. We thank the Bordeaux Imaging Center (BIC, a service unit of the University of Bordeaux and a national infrastructure, France BioImaging) where microscopy was performed and partially analyzed. This work was supported by ANR (NotifX, no. ANR-20-CE16-0016, to CM) and FRM (FDT202404018556, to SL).

We dedicate our article to the memory of Isabelle Caillé.

## Author contributions

Simon Lecomte: Designed research; Performed research; Analyzed data; Curated data; Wrote the first draft of the paper; Edited the paper; Wrote the paper, Acquired funding (FRM).

Bruna Ferreira Mira: Performed research; Analyzed data. Noelle Grosjean: Performed research; Analyzed data.

Christophe Mulle: Designed research; Supervised research: Edited the paper; Wrote the paper; Acquired funding (ANR).

---

**Figure S1.**
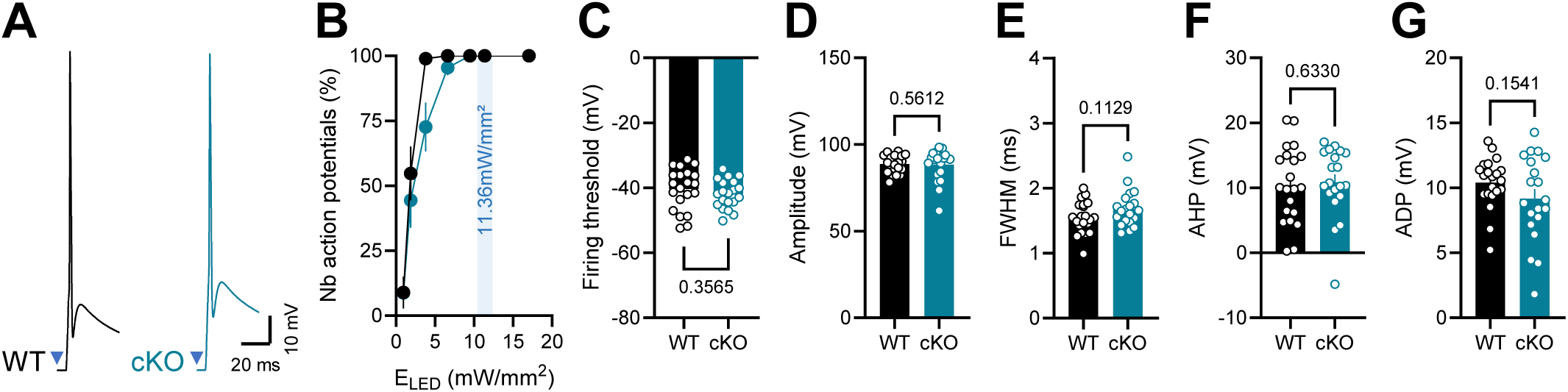
(related to figure 2). Lack of FMRP does not impact the properties of optogenetically-induced somatic action potentials from dentate granule cells. (A) Representative action potentials recorded after 1 ms light pulse of 470 nm at 11.36 mW/mm² from dentate granule cells in acute brain slices from control (black) and FMRP DG-cKO (blue) mice. Action potentials were elicited at 1 Hz. (B) Number of action potentials as a function of LED irradiance (ELED) at the focal plane. At 11.36 mW/mm² the stimulation is reliable enough to always elicit action potentials in the soma of granule cells in control and FMRP DG-cKO mice (blue area). Thus, 11.36 mW/mm² has been chosen for action potential properties comparison analysis. (C) The firing threshold is not affected by the lack of FMRP (Welch’s t-test, *t*(35.4)=0.9, *p*=0.3565). (D) The amplitude of action potential is not affected in the absence of FMRP (Mann-Whitney test, *U*=187, *p*=0.5612). (E) The full-width at half maximum is not changed by the lack of FMRP (Welch’s t-test, *t*(37.6)=1.6, *p*=0.1129). (F) The afterhyperpolarization phase of the action potential is not altered by the lack of FMRP (Mann-Whitney test, *U*=191, *p*=0.633). (G) The afterdepolarization phase of the action potential is not altered by the lack of FMRP (Welch’s t-test, *t*(30.4)=1.5, *p*=0.1541). WT: *n*=21 cells; cKO: *n*=20 cells. WT: *N*=7 mice; cKO: *N*=6 mice. Data are shown as mean ± SEM.

**Figure S2.**
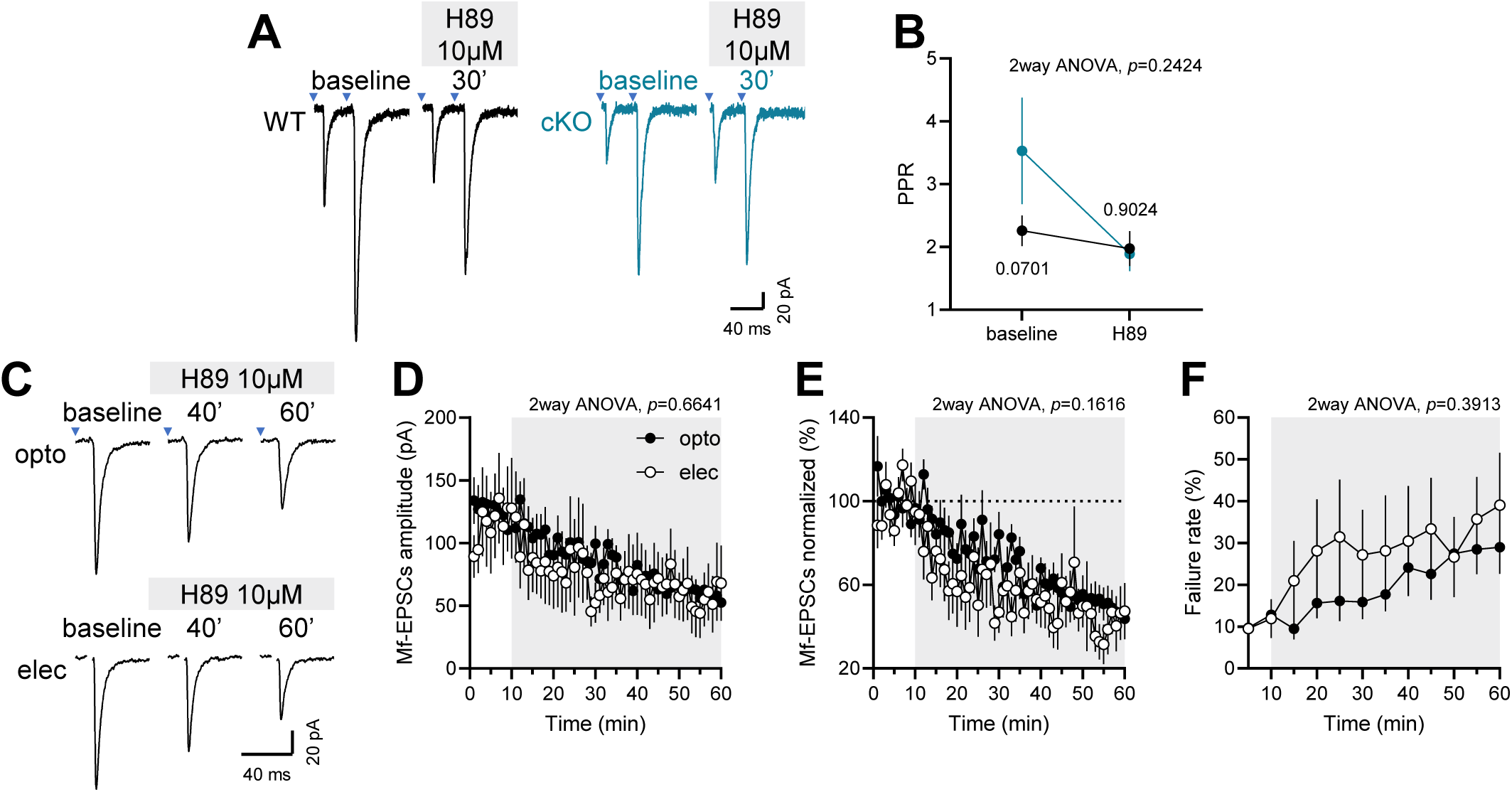
(related to figure 7). Presynaptic FMRP deletion does not affect release probability during H89 application, and H89 produces similar inhibition of electrically and optogenetically evoked Mf-EPSCs. (A) Representative traces of Mf-EPSCs obtained during a paired-pulses protocol in acute brain slices of control (black) and FMRP DG-cKO (blue) mice. Paired-pulses recordings (10 times) have been performed before and after 30 min H89 (10 µM) application. (B) Paired-pulses ratios before and after H89 do not differ between control and FMRP DG-cKO mice (2way ANOVA, main effect between groups, *F*(1, 18)=1.5, *p*=0.2424), neither during baseline (Fisher’s LSD post-hoc test, *p*=0.0701) nor after H89 application (Fisher’s LSD post-hoc test, *p*=0.9024). WT: *n*=10 cells; cKO: *n*=10 cells. WT: *N*=10 mice; cKO: *N*=8 mice. (C) Representative traces of Mf-EPSCs recorded at 0.1 Hz during 10 min baseline, 30 min and 50 min after H89 application (10 µM, grey area). Recordings from CA3 pyramidal neurons in acute brain slices from control mice during optogenetic or electrical stimulation. (D) Time course of Mf-EPSCs amplitude during 10 min baseline and during 50 min H89 (10 µM) application shows similar amplitudes between optogenetic (black circles) and electrical (open circles) stimulation (2way ANOVA, main effect between groups, F(1, 18)=0.2, *p*=0.6641). Time points are made by doing the average amplitude of 6 Mf-EPSCs. (E) Time course of normalized Mf-EPSCs amplitude during 10 min baseline and during 50 min H89 (10 µM) application shows similar amplitudes between optogenetic and electrical stimulation (2way ANOVA, main effect between groups, *F*(1, 18)=2.1, *p*=0.1616). Time points are made by doing the average amplitude of 6 Mf-EPSCs. (F) Time course of failure rate of Mf-EPSCs during 10 min baseline and during 50 min H89 (10 µM) application is not statistically different between optogenetic and electrical stimulation (2way ANOVA, main effect between groups, *F*(1, 18)=0.8, *p*=0.3913). Time points are made by calculating failure rate during 5 min of recording. opto: *n*=13 cells; elec: *n*=7 cells. opto: *N*=11 mice; elec: *N*=6 mice. Data are shown as mean ± SEM.

